# Gammaherpesvirus ncRNAs share conserved features of binding and virulence despite lack of sequence conservation

**DOI:** 10.1101/2022.05.09.491269

**Authors:** Ashley N. Knox, Eva M. Medina, Darby G. Oldenburg, Eric T. Clambey, Linda F. van Dyk

## Abstract

Gammaherpesvirus (γHV) non-coding RNAs (ncRNAs) are integral modulators of viral infection. The γHVs engage RNA polymerase III (pol III)-dependent transcription of both host and viral ncRNAs, which contribute to viral establishment, gene expression, and pathogenesis. Viral ncRNAs, such as the EBV-encoded RNAs (EBERs), reportedly interact with multiple host RNA-binding proteins (RBPs) and contribute to inflammatory responses implicated in the development of malignancies. Here, we examined RBP interactions of the pol III-transcribed tRNA-miRNA encoded non-coding RNAs (TMERs) of murine γHV68, and the potential contributions of these and the related EBERs to *in vivo* pathogenesis. Using sequential enzymatic treatments, we found that several TMER1 forms retain a 5’-triphosphate, lending the possibility of recognition by the innate immune sensor RIG-I. We further examined the interactions of TMERs and EBERs with host RBPs, and found that multiple TMERs and EBERs interact with the La protein, though minimal interaction was detected with RIG-I during primary virus infection. Finally, we investigated the contributions of the TMERs and EBERs to disease in an immune-compromised mouse model with a series of viral recombinants. We found that expression of multiple single TMERs, or the EBERs expressed in place of the TMERs, was capable of restoring virulence to a viral recombinant lacking expression of all TMERs. Ultimately, these studies demonstrate that divergent pol III-transcribed γHV ncRNAs share interaction characteristics with two host RBPs and conserved contributions to disease, despite little to no significant sequence conservation. These findings support a model of convergent functions of the sequence-variable pol III-transcribed γHV ncRNAs.

**IMPORTANCE:** Viruses manipulate the infected cell and host inflammatory responses through expression of coding and non-coding RNAs. The gammaherpesviruses are a subfamily of herpesviruses associated with chronic inflammatory diseases and malignancies, especially in immune-compromised individuals. Among these, the human Epstein-Barr virus and murine gammaherpesvirus 68 (γHV68) express highly abundant, RNA polymerase III-dependent, short non-coding RNAs. Whether these sequence-divergent ncRNAs have conserved functional properties is unknown. By using viral recombinants to allow direct comparison of these ncRNAs during primary infection, we find that the sequence-divergent ncRNAs of γHV68 and EBV share a conserved property to bind to the host RNA binding protein, La, and function interchangeably to facilitate *in vivo* pathogenesis. These studies demonstrate that abundant, RNA polymerase III-dependent viral ncRNAs can potently function to alter the host cell landscape and promote disease in a sequence-independent manner.

## INTRODUCTION

The gammaherpesviruses (γHVs) are DNA viruses that establish life-long infection in a lymphocyte reservoir within their hosts (1, 2). The human-specific γHVs include Kaposi’s sarcoma associated herpesvirus (KSHV or HHV-8) and Epstein-Barr virus (EBV or HHV-4). Murine gammaherpesvirus 68 (MHV68 or γHV68; ICTV nomenclature *Murid herpesvirus 4*, MuHV-4) serves as a small animal model of γHV infection and pathogenesis (3, 4). Primary infection with the γHVs is characterized by high viral gene expression and production of new virions. Following resolution of this lytic phase, the virus is maintained in a latent state, during which viral gene expression is low and new virions are not produced. Latency is maintained through the establishment of an equilibrium between the virus and the host immune system in healthy immune-competent hosts; however, disruption of this balance is associated with virus reactivation, leading to a range of malignancies (5).

The γHVs express several types of non-coding RNAs (ncRNAs) that play important roles in the virus lifecycle. Interestingly, some γHV ncRNAs are transcribed by RNA polymerase III (pol III), lending them unique transcriptional regulation and characteristics. The γHV68 tRNA- miRNA-encoded RNAs (TMERs; TMER1-TMER8) are expressed during lytic and latent infection (6). The TMERs are dispensable for lytic replication and the establishment of latency, but required for pathogenesis (7, 8). The EBV-encoded small RNAs (EBERs; EBER1 and EBER2) are also expressed during both lytic and latent infection and have been shown to interact with several host proteins, including ribosomal protein L22, protein kinase R (PKR), lupus-associated antigen (La), and retinoic acid-inducible gene I (RIG-I) (9). Interactions between EBERs and host proteins can trigger sustained host innate immune responses that are implicated in the development of EBV-associated malignancies (10–13), highlighting an integral role of pol III-transcribed RNAs in γHV pathogenesis.

Certain host RNA-binding proteins (RBPs) are predicted to bind the EBV EBERs through features that are necessarily imparted by pol III transcription (14). Unlike RNA pol II- transcribed RNAs, pol III transcripts are not modified to add a 5’ 7-methylguanine cap, leaving a 5’-triphosphate on pol III-transcribed primary RNAs (14, 15). Pol III transcription is terminated by a stretch of thymine residues, transcribed to a 3’-polyU sequence (16, 17). Additionally, the predicted secondary structures of EBER1 and EBER2 include sections of double-stranded RNA (dsRNA) (18). The innate immune sensor RIG-I binds to RNAs with a 5’-triphosphate and dsRNA, and the La protein binds 3’-polyU sequences characteristic of all pol III-transcribed primary RNAs (16, 17, 19, 20). EBER interaction with RIG-I reportedly induces type I interferon and IL-10 signaling, while binding of EBERs with La initially facilitated EBER discovery and putatively mediates interaction with TLR3, leading to type I interferon and inflammatory cytokine responses (10-13, 21, 22). Due to the strict human-specificity of EBV, these studies largely rely on the analysis of patient samples or the comparison of established EBV-negative and EBV-positive transformed cell lines that are latently infected. Therefore, the understanding of EBER interactions with host proteins during primary lytic infection is limited.

Like the EBV EBERs, the γHV68 TMERs are transcribed by pol III (23). These bifunctional RNAs contain a 5’ tRNA-like structure followed by multiple hairpins that are processed into biologically active miRNAs (8, 24–26). Sequence-dependent targets of the TMER miRNAs are highly conserved with KSHV and EBV, and include pathways in host translation and protein modification (27). Our previous work characterizing a γHV68 recombinant lacking expression of all eight TMERs (TMER total knock-out, TMER-TKO γHV68) demonstrated the dispensability of TMERs for lytic replication, while highlighting the requirement of TMERs for optimal virulence in an acute pneumonia model (7). Furthermore, expression of the tRNA-like portion of TMER1, in the absence of any γHV68 miRNAs, restored pathogenesis. Our data suggest that the tRNA-like sections of the TMERs act as sequence-independent modulators of infection, potentially through interactions with host RBPs.

Due to the similarities between the EBERs and TMERs (pol III transcription and predicted dsRNA secondary structure), host RBP interactions with EBERs imply similar interactions with TMERs. However, unlike EBERs, TMERs undergo extensive processing that result in multiple forms that may differ in their capacity as ligands for host RBPs. The tRNA-like portion of the TMERs has been shown to be processed into a mature, non-aminoacylated tRNA (24), and TMERs contain alternate transcription terminators and hairpins that are processed into miRNAs (8, 28). Whether the various forms of TMERs maintain characteristics that are recognized by host RBPs like RIG-I and La remains unknown. Our previous work using reverse ligation mediated reverse transcription PCR (RLM RT-PCR) indicates the presence of 5’- monophosphate on some mature TMER miRNAs (8), however, characterization of the 5’-end of the primary TMERs has not yet been reported.

Identifying interactions between γHV ncRNAs and host RBPs is integral to understanding the role of γHV ncRNAs in host immune modulation and pathogenesis. By analyzing the modifications on the γHV68 TMERs, we found evidence that some species retain a 5’-triphosphate end, indicating the potential of these RNAs to bind to RIG-I. Subsequent studies found that the TMERs and EBERs bound to multiple RBPs with varying stringency, with a particular robust interaction observed with the La RBP. Our data further indicate that while the EBERs and TMERs lack sequence conservation, both classes of γHV ncRNAs share the capacity to enhance *in vivo* pathogenesis in an acute model of γHV disease. Ultimately, our studies show that the pol III-transcribed γHV ncRNAs lack sequence conservation, but have shared binding characteristics with two host RBPs and drive pathogenesis.

## RESULTS

### 5’-end characterization demonstrates γHV68 TMERs with 5’-triphosphate ends

To investigate potential interactions of the γHV68 ncRNAs with the innate immune sensor, RIG-I, we sought to characterize the 5’ RNA ends using a previously published sequential enzymatic assay (29). Small RNAs were isolated and size fractionated from HEK 293 cells 24 hours after mock treatment or after infection with WT or TMER1-only γHV68. The TMER1-only γHV68 was previously characterized, and only expresses TMER1 but no other TMERs, allowing for specific analysis of an individual TMER (7). All RNAs smaller than 300 nucleotides (Fig 1A) were subjected to three different enzyme treatments; 1) RNA 5’- polyphosphatase only, 2) Terminator™ only, or 3) RNA 5’-polyphosphatase followed by Terminator™ (Fig 1B). Untreated RNA was included as a control. RNA 5’-polyphosphatase converts 5’-triphosphate to 5’-monophosphate RNAs, and Terminator™ is a 5’-monophosphate-dependent exonuclease. Therefore, we expect any RNAs with a 5’-triphosphate to resist degradation when treated only with Terminator™ enzyme (identified as “-/+”, Fig 1B), but to be sensitized to Terminator™ degradation when first treated with RNA 5’-polyphosphatase (“+/+”). These two populations are highlighted in blue boxes throughout Figures 1 and 2 to indicate which populations were compared to detect the presence of 5’-triphosphate RNAs.

**Figure 1.**
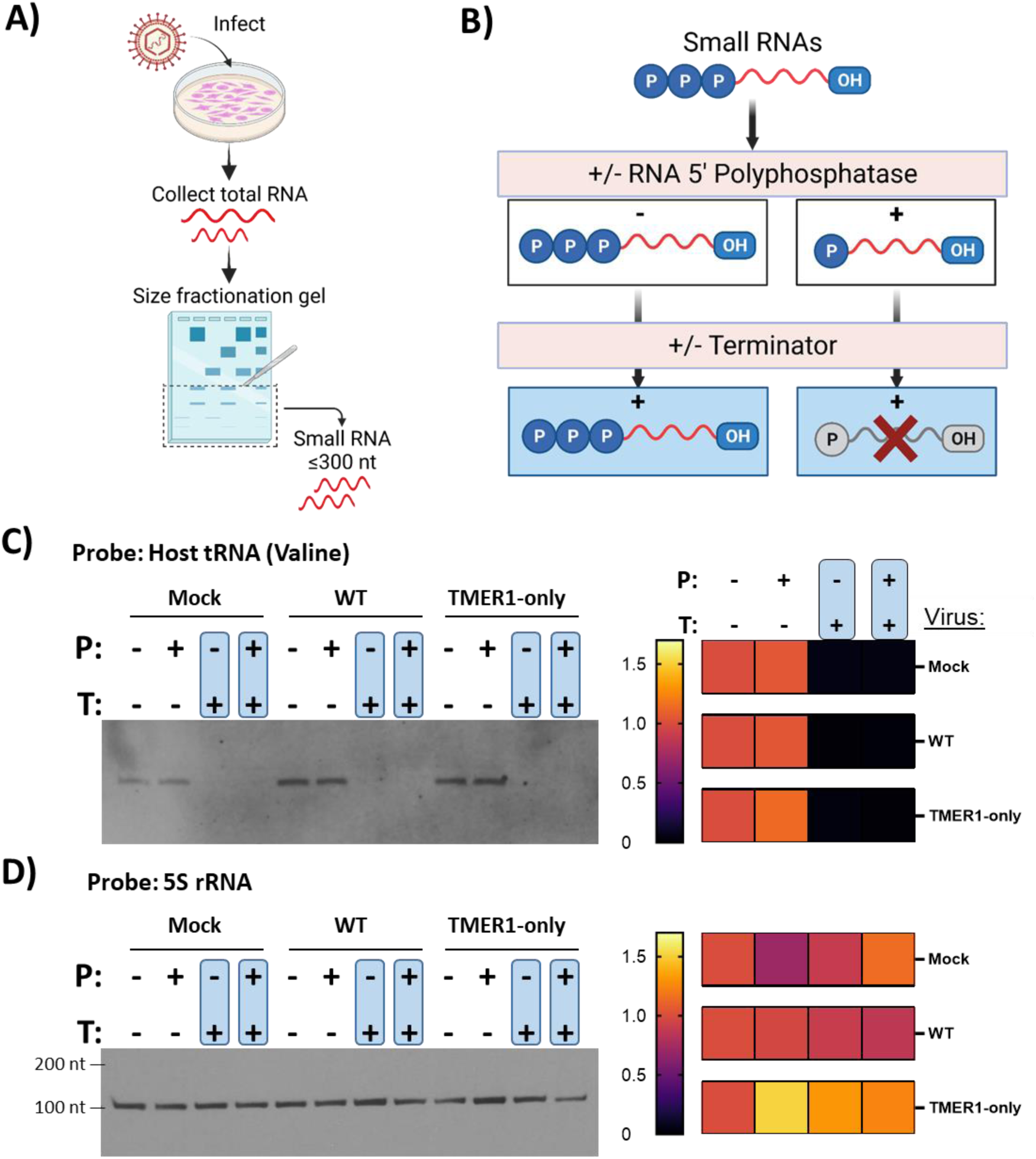
Sequential enzyme treatments distinguish between susceptible and resistant 5’ RNA ends. **A)** Experimental design schematic for isolating small RNAs. HEK 293 cells +/- infection with WT or TMER1-only γHV68 were incubated for 24 hpi prior to total RNA collection and size fractionation for small RNAs under 300 nts. **B)** Experimental design schematic for 5’ end characterization of RNA. Small RNAs are treated with or without RNA 5’-polyphosphatase to convert 5’-triphosphate to 5’-monophosphate ends, and then treated with or without Terminator™ enzyme to degrade only RNAs with a 5’-monophosphate. Human tRNA valine **(C)** and human 5S rRNA **(D)** 5’ end characterization: enzymatic products indicated were resolved by SDS-PAGE and northern blots tested with probes specific to host tRNA or 5S rRNA transcripts (left). Densities of the resulting bands were normalized to an ethidium bromide stained 5S rRNA loading control. Relative band density was calculated as a fold change of the untreated RNA population, which was set to 1, and displayed as heat maps (right). Heat maps represent n = 4 for tRNA and n = 3 for 5S rRNA.

**Figure 2.**
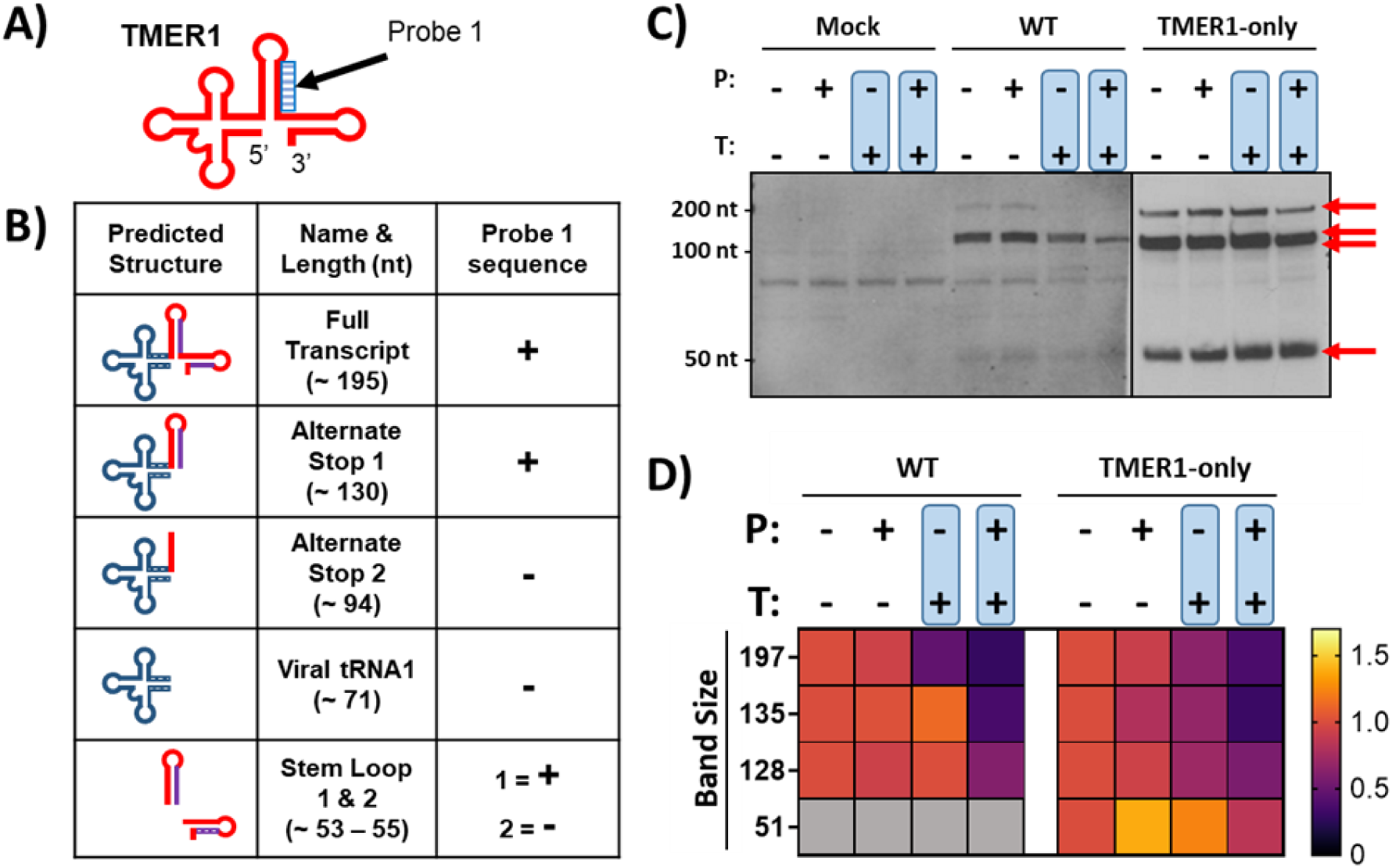
RNA end characterization indicates TMER1 derived RNAs with 5’-triphosphate ends. **A)** Schematic of the predicted secondary structure of TMER1 with sequence for northern TMER1 probe 1 indicated at the 3’ side of stem loop 1. **B)** TMER1 primary and processed forms: predicted structures in left column, name and length in center, and TMER1 probe 1 sequence in right column (“+” indicates sequence present; “-” indicates sequence not contained). The tRNA-like loop is shown in blue, and TMER1 miRNAs in purple. **C)** Following sequential enzymatic treatments as detailed in Figure 1B, small RNAs were resolved by SDS-PAGE then detected by northern blot with TMER1 probe 1. Blot is representative of three independent experiments. RNA bands detected only during infection and specific to TMER1 are marked with red arrows, in contrast to non-specific bands shared with mock infected samples. **D)** Densities of the TMER1 northern blot bands were normalized to an ethidium bromide stained 5S rRNA loading control. Relative density of bands were calculated as a fold change of the untreated RNA population, which was set to 1, and presented as heat maps. Band sizes were calculated as averages based on migration of a ladder included in each experiment. RNA bands not detectable in WT γHV68 infection are shown as gray boxes. Data is from three independent experiments.

We first characterized the human (host) tRNA-Valine and 5S rRNA (Fig 1C-D) following sequential enzyme treatment. Specific RNAs were detected by northern blot probes (Table 1) and the density of each population was normalized to an ethidium bromide-stained 5S rRNA control. To determine how each RNA responded to enzyme treatment, the band density of each enzyme-treated population was calculated as the fold change of the untreated population (-/-), which was set to 1. Fold changes are presented as heat maps (Fig 1C-D). Our analysis indicates complete degradation of the host tRNA following treatment with Terminator™ (Fig 1B-C), as expected due to rapid processing of tRNA 5’ ends by RNase P (reviewed in (30)). Interestingly, there was no significant degradation of the human 5S rRNA following both enzyme treatments, despite a well characterized 5’-triphosphate end and a previous report showing significant 5S rRNA degradation following both enzyme treatments (29). The difference in our analysis may be due to lower sensitivity of the assay, and considering that 5S rRNA is a highly abundant transcript, any degradation in this population may fall below our limit of detection.

**Table 1.**
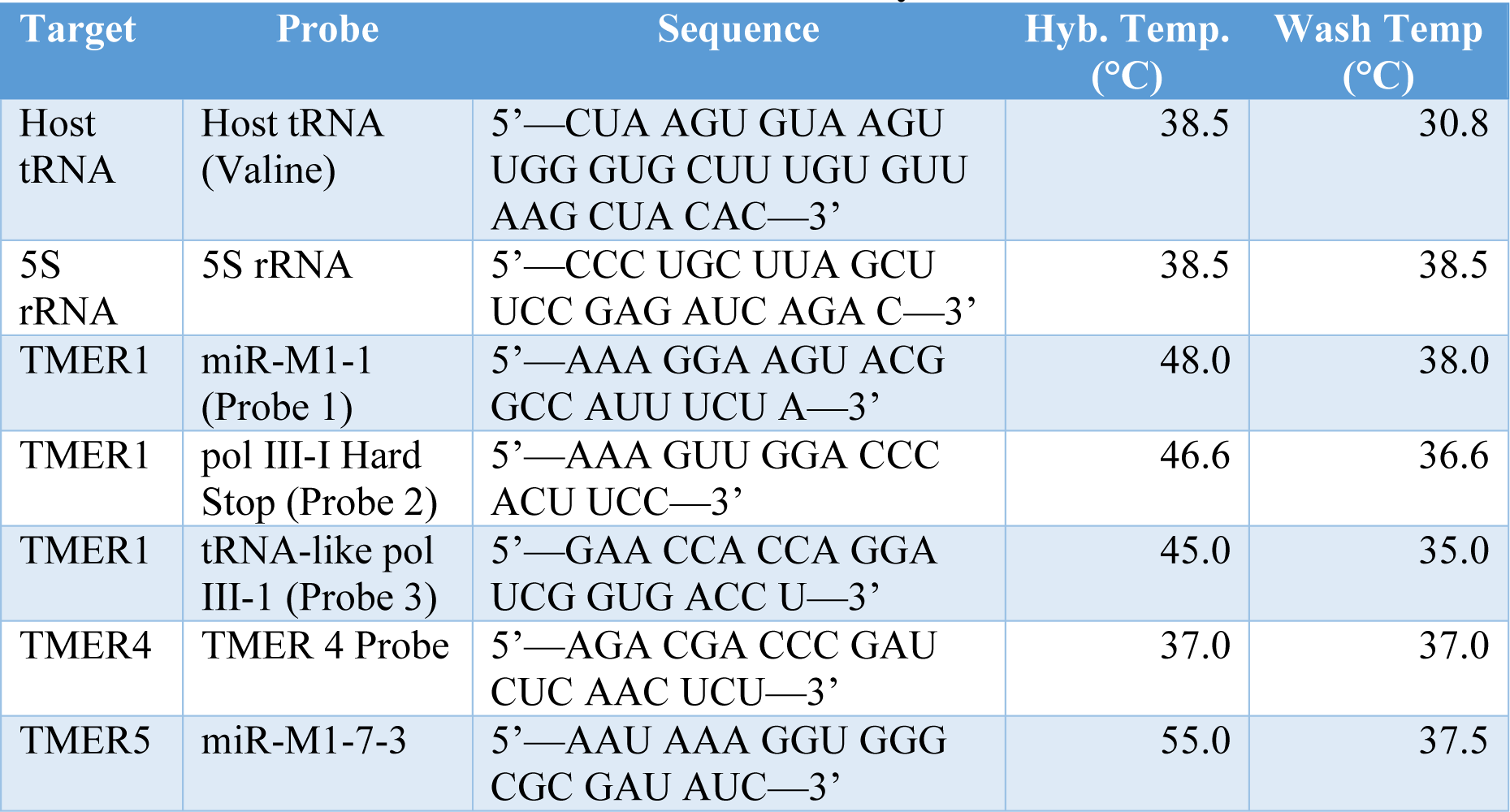
Probes and conditions for northern blot analyses.

We then used sequential enzymatic treatment to characterize the 5’ end of the most abundant γHV68 TMER, TMER1. The predicted secondary structure of TMER1 RNA is shown in Fig 2A with the location of the northern probe indicated (Probe 1). The TMER1 gene contains two alternative transcriptional termination sites, and can be processed into a tRNA-like 5’ portion and multiple hairpins that give rise to biologically active miRNAs (8, 24, 28). These characteristics result in multiple possible forms of TMER1 RNA with various lengths, some of which are detected with Probe 1 as indicated (Fig 2B). Following sequential enzyme treatment as previously described, TMER1 RNA was detected by northern blot (Fig 2C). Mock-treated samples indicate some off-target bands, while several RNA populations were only detected in the WT and TMER1-only γHV68 infected samples specific to TMER1 (Fig 2C, red arrows). As expected, we detected TMER1 RNA at multiple sizes consistent with known alternative forms of TMER1. We previously observed that TMER1 is expressed more abundantly during infection with TMER1-only recombinant virus compared to WT virus (7), as observed here where TMER1 RNA is more abundant in TMER1-only than WT γHV68-infected samples. Density analysis was performed on each band (standardized using an RNA ladder) and presented as the fold change compared to the untreated RNA population as an average across three independent experiments (Fig 2D). Notably, several TMER1 populations displayed increased Terminator™ degradation following pretreatment with RNA 5’-polyphosphatase, indicating the presence of 5’- triphosphates on some TMER1 RNAs (e.g. the ∼195 nt species of TMER1, Fig 2D). Analysis of TMER1 RNAs with other northern probes (Fig S1) provides further evidence of 5’-triphosphate containing TMER1 RNAs (e.g. the ∼195 nt band, Fig S1D-G).

The γHV68 TMERs show conservation of predicted secondary structures, and therefore we extended our 5’ characterization analysis to determine whether 5’ ends differ or are consistent across multiple TMERs. As with TMER1, northern probe sequences in TMER4 or TMER5 bind multiple processed forms, resulting in several band sizes detected by northern blot. Band density analysis did not indicate the presence of TMER4 or TMER5 species maintaining a 5’- triphosphates (Fig S2). Our characterization of TMER1 RNA benefited from the use of the TMER1-only recombinant virus, which resulted in more abundant TMER1 RNA and more sensitive TMER1 detection (Fig 2C northern blots from TMER1-only infection vs WT infection). However, our characterization of TMER4 and TMER5 relied solely on WT γHV68- infected samples, as the TMER4 and TMER5 only recombinants were not yet available at the time of this analysis. Future studies with additional γHV68 recombinants may facilitate more detailed 5’ end characterization of these TMERs.

### TMER and EBER ncRNAs are detected in RIG-I and La immunoprecipitated complexes under permissive conditions

Detection of 5’-triphosphate TMERs suggests these RNAs are potential ligands for host retinoic acid inducible gene I (RIG-I), a pattern recognition receptor previously reported to bind double-stranded RNA with 5’-triphosphates to induce innate immune pathways (19, 20). The TMERs and EBERs are transcribed by RNA polymerase III (pol III), and therefore end with 3’- oligouridylate (3’-poly(U)) for potential binding to host La protein (16, 17). Considering these characteristics of the γHV68 TMERs and previous EBER reports, we investigated whether they interact with the host RBPs RIG-I and La.

We transfected HEK 293 cells with plasmids expressing either FLAG-tagged RIG-I or La (Fig 3A). One set of samples were also transfected with an EBER-expressing plasmid to compare the different γHV ncRNAs. Following transfection, cells were infected with either the TMER1-only γHV68 or a previously characterized viral recombinant that does not express any of the TMERs (TMER-total knockout, TMER-TKO γHV68). The EBER-plasmid transfected cells were infected with the TMER-TKO γHV68 to provide shared infection conditions with only EBER ncRNAs. Cell lysates were collected 24 hpi and immunoprecipitated with FLAG-specific magnetic beads. To maximally detect viral ncRNAs potentially bound to host RBPs, we first examined permissive conditions, in which the immunoprecipitates were washed three times with TBS. A fraction of the IP complexes were analyzed by western and remaining beads were subjected to TRIzol for RNA isolation. Western blots confirmed the effective purification of the intended host proteins (Fig 3B). Isolated RNA analyzed by reverse transcription polymerase chain reaction (RT-PCR) with primers targeting either TMER1 or EBER1 demonstrated that both γHV ncRNAs were detected in the RIG-I and La immunoprecipitates under conditions of low stringency washes and high cycle number (Fig 3C). We extended this analysis to another highly expressed γHV68 ncRNA, TMER5. HEK 293 cells were treated as before, but were also infected cells with the WT γHV68, which expresses all TMERs. We found that TMER5 was also detected in these host RBP immunoprecipitates with low-stringency washes and high cycle number (Fig S3A-B). Together, these studies suggested that the γHV TMERs and EBERs are alike in their relative binding to the RBPs La and RIG-I during primary infection.

**Figure 3.**
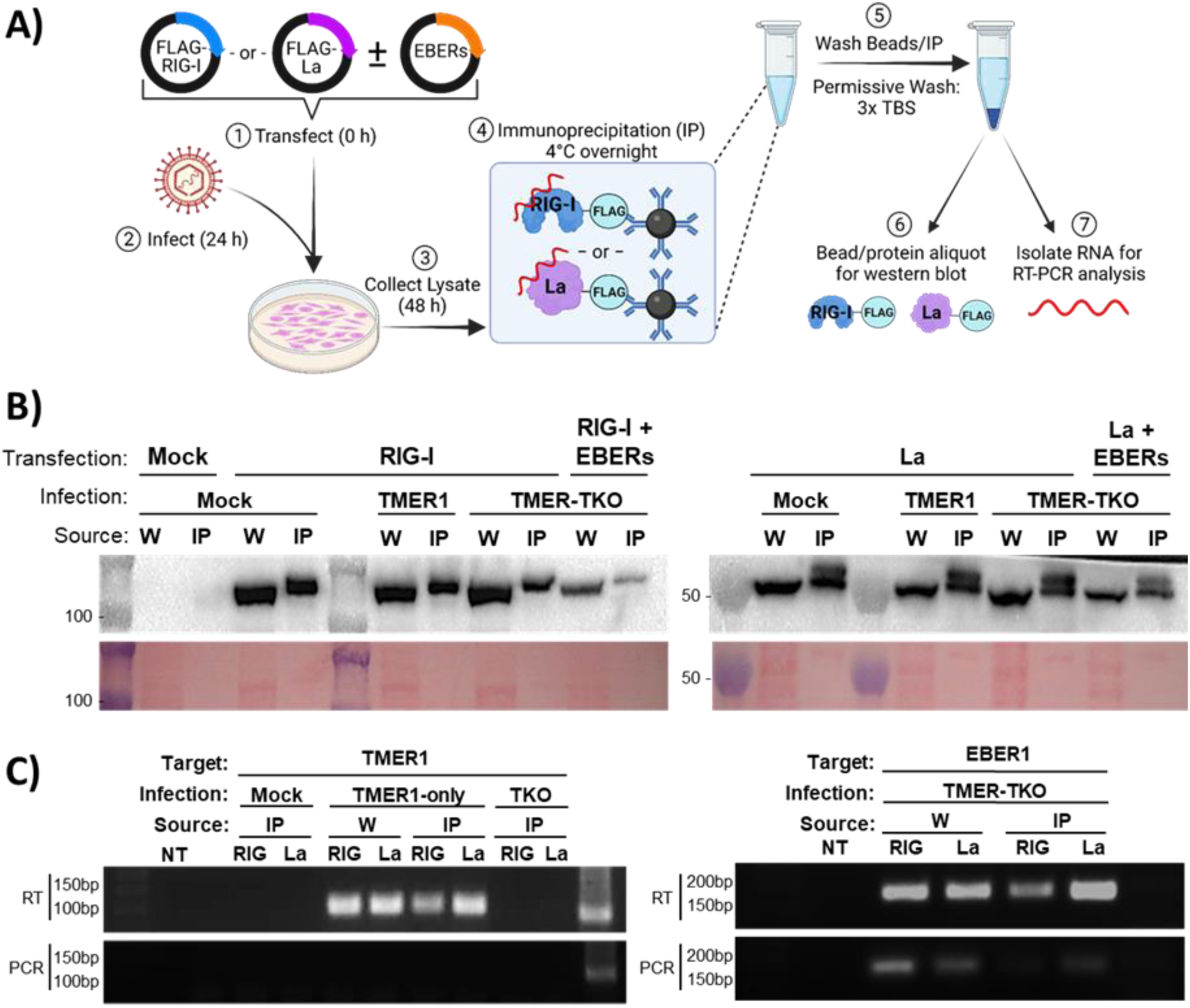
FLAG-mediated immunoprecipitation of RNA binding proteins and associated RNAs under permissive conditions. **A)** Experimental design to study ncRNA interactions with RIG-I and La. HEK 293 cells were transfected with FLAG-tagged proteins of interest; RIG-I or La. For EBER interaction analysis, cells were also transfected with a plasmid expressing both EBER1 and EBER2 (pSP73-EBERs). 24 hours after transfection, cells were infected with mock, WT, TMER1-only, or TMER-TKO γHV68 at an MOI of 5. 24 hpi (48 hours post-transfection), immunoprecipitation was performed, followed by “permissive wash” with TBS. An aliquot of beads was reserved for western blot analysis (B) and RNA was isolated from the remaining beads for RT-PCR (C). **B)** Proteins from whole cell lysate (W) or immunoprecipitation beads (IP) were resolved by SDS-PAGE and detected by western blot with a primary antibody targeting FLAG for RIG-I-FLAG (left) or La- FLAG (right) transfected samples. Ladder shows protein size in kDa. Ponceau red stained blots, below, demonstrate enrichment by IP. Blots are representative of two independent experiments with technical triplicates. **C)** RNA was isolated from whole cell lysate (W) or immunoprecipitated (IP) samples from cells transfected with RIG-I-FLAG (RIG) or La-FLAG (La). Primers targeting TMER1 (left) or EBER1 (right) were used for RT-PCR with 40 cycles. PCR without reverse transcription (“PCR”) was performed in conjunction with RT-PCR to test for DNA contamination. Data are representative of two independent experiments with technical duplicates or triplicates.

### TMER and EBER ncRNA bind to La, but not RIG-I, under stringent conditions

Though we were able to detect multiple TMERs and EBER1 in RIG-I and La immunoprecipitates, we wanted to further examine the specificity and strength of these interactions in a highly stringent, quantitative analysis. Therefore, we modified the immunoprecipitation procedure to include a control protein not known to bind RNA (FLAG-tagged GFP) and stringent wash conditions following immunoprecipitation (Fig 4A).

**Figure 4.**
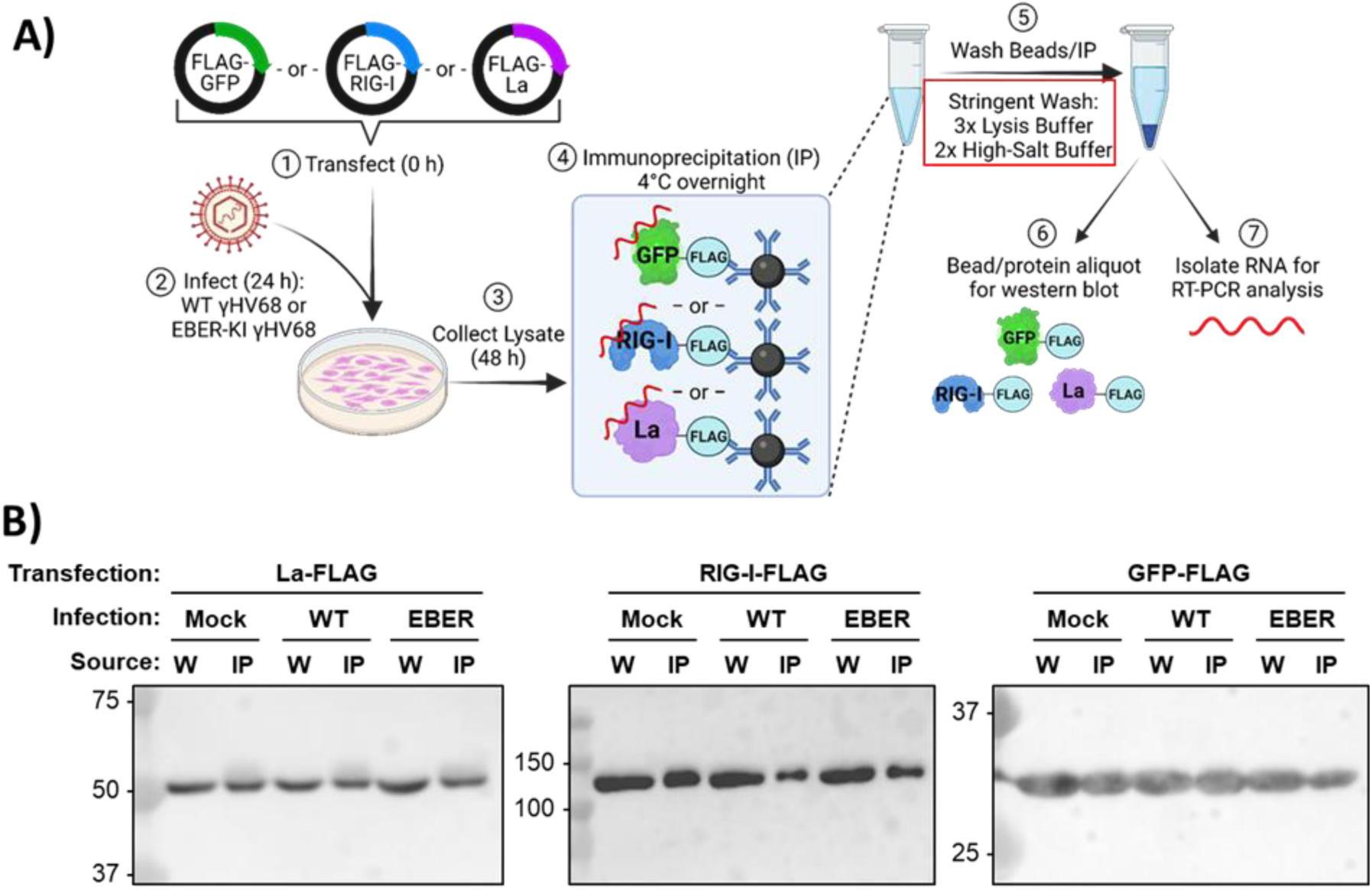
Modified immunoprecipitation experiment with a more stringent wash successfully isolates FLAG-tagged proteins of interest. **A)** Modified experimental design to detect ncRNA interactions with RIG-I and La. Experiment was performed as previously described (Figure 3) with the following modifications. HEK 293 cells were transfected with FLAG-tagged RIG-I, La, or GFP as a non-specific binding control. 24 hours after transfection, cells were infected with WT or EBER-knock in (EBER-KI) γHV68. Immunoprecipitation was performed as before, followed with a “stringent” wash of beads (outlined in red box) prior to protein analysis and RNA isolation as before. **B)** Proteins from whole cell lysates (W) or immunoprecipitated beads (IP) were resolved by SDS-PAGE and western blot analysis was performed with a primary antibody targeting FLAG. Proteins were analyzed in mock, WT γHV68 infection (WT), or EBER-KI γHV68 infection (EBER). Protein ladder is indicated to the left of each blot (kDa). Expected approximate protein sizes: La = 47 kDa, RIG-I = 102 kDa, GFP = 27 kDa.

Additionally, we used a new γHV68 recombinant that expresses both EBV EBERs in place of the TMERs. The EBER-knock in (EBER-KI) βla.γHV68 virus expresses both EBERs in place of the TMERs but maintains the rest of the γHV68 genome, allowing us to examine the function of EBERs in *de novo* γHV infection. We previously reported that EBER-KI βla.γHV68 infects murine fibroblast cells and expresses EBERs without any detectable TMER expression as measured by RNA-based flow cytometry (31). With this tool established, we infected HEK 293 cells with WT or EBER-KI βla.γHV68 and performed immunoprecipitation of FLAG-tagged proteins as before, but following IP, bead-protein complexes were subjected to a stringent wash step consisting of three washes with lysis buffer followed by two washes with a high-salt buffer (Fig 4A, red box), based on previously published wash conditions for RIG-I bound RNAs during HSV-1 infection (32). Samples were analyzed by western blot to confirm effective immunoprecipitation (Fig 4B).

RNA isolated from immunoprecipitates was initially subjected to RT-PCR with primers targeting TMER1 or EBER1 (Fig 5A-B) and a limited number of RT-PCR cycles. We reasoned that meaningful interaction between the γHV ncRNAs and the RBPs of interest would result in an enriched signal in IP samples compared to total RNA samples, and non-specific interactions would fall below the limit of detection. Under these conditions, we detected strong interactions of both TMER1 (Fig 5A) and EBER1 (Fig 5B) with the La protein; however, both γHV ncRNAs demonstrated signal with FLAG-RIG-I that was at or below the signal observed with the FLAG-GFP negative control. To determine if these observations were common across other γHV ncRNAs, we repeated semi-quantitative RT-PCR analysis with primers targeting TMER4 (Fig 5C) and EBER2 (Fig 5D). We found that TMER4 and EBER2 were strongly detected from La but not RIG-I samples, suggesting that both TMERs and EBERs consistently interact with the La protein during *de novo* primary γHV infection (Fig 5C-D).

**Figure 5.**
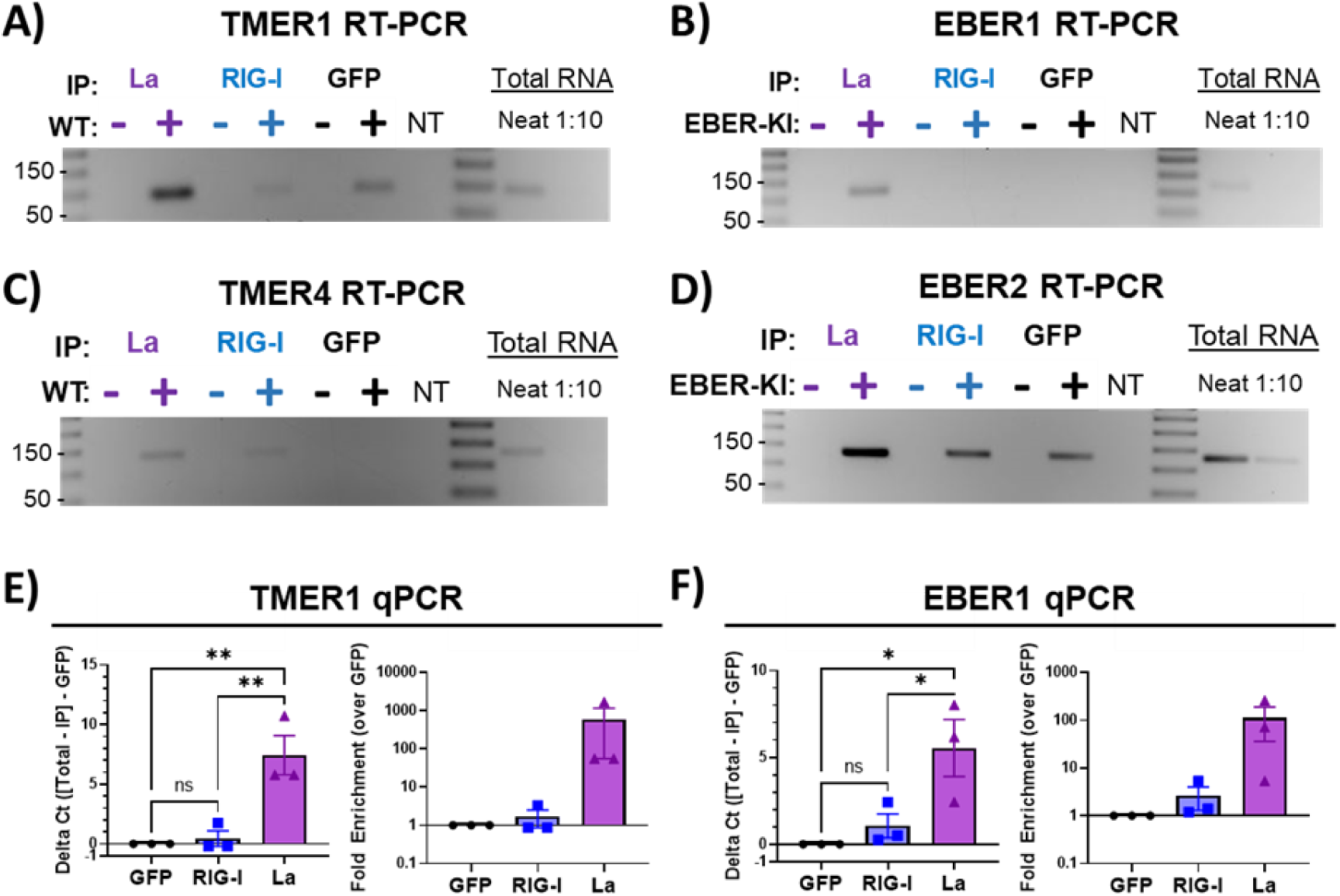
FLAG-mediated immunoprecipitation of RNA binding proteins and associated RNAs under stringent conditions. RNA was isolated from immunoprecipitation complexes as previously described. RT-PCR for TMER1 **(A),** EBER1 **(B),** TMER4 **(C)**, and EBER2 **(D)** was limited to 30 cycles or 33 cycles (TMER4) to detect enrichment of the target RNA in IP samples compared to a neat and diluted total RNA positive control. Numbers to the left of each gel indicate ladder sizes (bp). Quantitative analysis was performed by RT-qPCR for TMER1 and EBER1. The ΔCt for TMER1 **(E)** and EBER1 **(F)** interacting with RIG-I or La was calculated as: (Total Target Ct – IP Target Ct) – (Total GFP Ct – IP GFP Ct), where “target” refers to RIG-I or La. ΔCt for GFP equals 0, while positive values indicate enrichment and negative values indicate diminishment of RNA interaction with target proteins, respectively. Fold enrichment for TMER1 **(E)** or EBER1 **(F)** was calculated as 2^𝛥𝐶𝑡^, where ncRNA detected with GFP is set to 1. Error bars = SEM. Significant differences analyzed by one-way ANOVA with multiple comparisons and indicated as asterisks. P-values are indicated as follows: * = P ≤ 0.05, ** = P ≤ 0.01.

To further quantify these interactions, we measured TMER1 and EBER1 using quantitative RT-PCR (RT-qPCR). RNA samples that were determined to be free of DNA contamination by comparable RT-PCR and PCR reactions were converted to cDNA, prior to TMER1 or EBER1 measurement by SYBR Green RT-PCR. The ΔCt values for TMER1 (Fig 5E) or EBER1 (Fig 5G) in GFP, RIG-I, and La samples were calculated by normalizing immunoprecipitated Ct values to total RNA Ct values, then subtracting the corresponding normalized Ct values for GFP samples. We found enrichment of both TMER1 and EBER1 in La samples compared to the negative control GFP samples; however, we did not detect significant enrichment in RIG-I samples (Fig 5E-F, left graphs). These observations were further corroborated when we then calculated the fold enrichment as 2^Δ𝐶𝑡^ for TMER1 and EBER1 (Fig 5E-F, right graphs). These data demonstrate that the γHV68 TMERs and EBV EBERs share a robust and specific interaction with the La protein during *de novo* primary infection.

### γHV68 ncRNA recombinants expressing individual TMERs or EBERs show normal replication *in vitro*

To better understand the specificity or redundancy of γHV ncRNAs, we next established a panel of γHV68 recombinants engineered to express individual TMERs or the EBERs. These γHV68 recombinants include TMER4-only, TMER5-only, TMER8-only, and the previously mentioned EBER-KI γHV68 (schematics in Fig 6A). Each recombinant was confirmed by restriction digestion and sequencing of the left end of the virus. The bacterial artificial chromosome (BAC) DNAs used to generate recombinants were confirmed by PCR analysis for expected correct ncRNA sequence (Fig 6B). Expression of the intended ncRNAs with these new recombinants was confirmed by RT-PCR of RNA isolated from infected HEK 293 cells (Fig 6C). Finally, the replication kinetics of each recombinant was compared to WT virus by single-step replication analysis, studies that demonstrated comparable *in vitro* replication between this series of recombinants (Fig 6D), indicating that the expression of any single TMER or the EBERs in place of the WT TMERs had a negligible impact on replication fitness *in vitro*. This observation is consistent with our previous report that the TMERs are dispensable for lytic replication (7).

**Figure 6.**
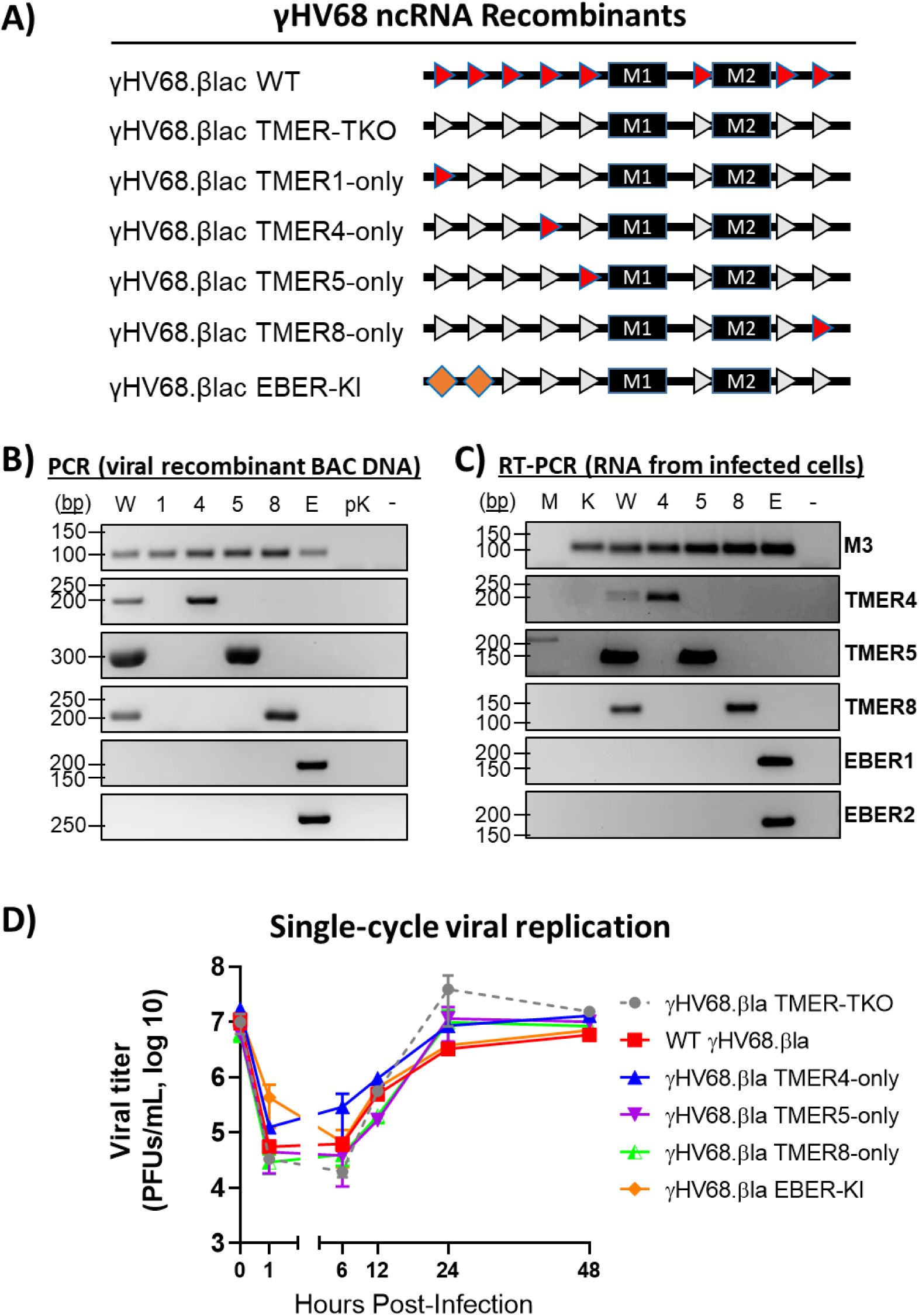
Characterization of γHV68 ncRNA recombinants. **A)** Schematics representing genetic details of the γHV68 ncRNA recombinants. Line diagrams represent the first 6 kilobases of the γHV68 genome, including the M1 and M2 genes (black rectangles). Each intact TMER gene is depicted as a red triangle. Gray triangles represent TMERs that are not expressed due to promoter deletion as previously described (7). Orange diamonds represent the expression of EBERs in place of TMERs through knock-in of the EBER1 and EBER2 sequences into the left end of the γHV68 genome (EBER knock-in; EBER-KI). **B)** PCR of viral recombinant DNA. W = WT γHV68, 1 = TMER1-only γHV68, 4 = TMER4-only γHV68, 5 = TMER5-only γHV68, 8 = TMER8-only γHV68, E = EBER-KI γHV68, pK = pLE—TMER-TKO plasmid as previously described ((7); does not contain M3), “-” = no-template control. Targets listed to the right of PCR panels. **C)** RT-PCR of RNA collected from HEK 293 cells infected with γHV68 recombinants at an MOI of 1. Viruses indicated as in B, except M = mock and K = TMER-TKO γHV68 (expresses M3). Targets for B and C listed to the right of PCR panels. PCR without reverse transcription was run with the same conditions as each RT- PCR to confirm the absence of DNA contamination (not shown). Some product sizes differ than the same target in (B) due to the use of different primers better suited to RT-PCR analysis. **D)** Single step replication analysis with WT γHV68 (red squares) or recombinants in 3T12 cells at an MOI of 5. Other viral recombinants shown are TMER-TKO (gray circles, dashed line), TMER4-only (blue triangles), TMER5-only (flipped purple triangles), TMER8-only (half-filled green triangles), and EBER-KI (orange diamonds). Cells and supernatants were collectively harvested at the indicated times post-infection, then quantified by plaque assay. Data depict the mean of 3 biologic replicates within a single experiment. Error bars = SEM.

### The γHV TMER and EBER ncRNAs share capacity for virulence *in vivo*

To study how the individual γHV ncRNAs recombinants replicate and contribute to virulence *in vivo*, we infected interferon gamma deficient (BALB.IFN-γ^-/-^) mice with the panel of viral recombinants and measured lung viral titer and survival. WT γHV68 has previously been demonstrated to cause acute pneumonia in BALB.IFN-γ^-/-^ mice, resulting in a high mortality rate by 14 days post-infection (33). However, mice inoculated with a γHV68 mutant with a 9,473-bp left-end deletion (which removes all eight TMERs, M1, M2, M3, and part of M4) display fully- attenuated pathogenesis, showing that viral genes in the left end of the genome are required for pathogenesis (33, 34). Studies of the BALB.IFN-γ^-/-^ model further showed that infection with the βla.γHV68 TMER-TKO results in reduced virulence compared to WT βla.γHV68, and expression of a single TMER (γHV68.βla TMER1-only) or the tRNA-like portion of TMER1 alone (γHV68.βla vtRNA1-only) reverses the pathogenic deficit of the TMER-TKO virus (7).

This previous study showed that the expression of only TMER1 is sufficient for virulence in the acute pneumonia mouse model. To determine if this pathogenic capacity is unique to TMER1 or a shared feature of γHV ncRNAs, we tested the new viral recombinants that express different single TMERs or both EBERs. BALB.IFN-γ^-/-^ mice were intranasally inoculated with each viral recombinant (4×10^5^ pfu/mouse) for analysis of *in vivo* replication from infected lung tissue 8 days p.i. by qPCR for viral DNA (gB sequence; Fig 7A) or by plaque assay (Fig 7B). We found no significant difference in *in vivo* replication among the viral recombinants, indicating equivalent replication *in vivo*. Inoculated BALB.IFN-γ^-/-^ mice were then monitored over the course of 15 days p.i. to determine morbidity in the pneumonia model (Fig 7C). The TMER-TKO βla.γHV68-infected mice showed significantly higher survival rate after 15 days than the WT βla.γHV68-infected mice, consistent with a previous report (7). Notably, all four of the new ncRNA βla.γHV68 recombinants showed no significant difference in pathogenesis compared to WT βla.γHV68. These results indicate that despite unique sequences across viral ncRNA genes, each individual viral ncRNA (TMER 4, 5, or 8) or the EBERs in place of the TMERs share the ability to promote *in vivo* virulence in an acute pneumonia model.

**Figure 7.**
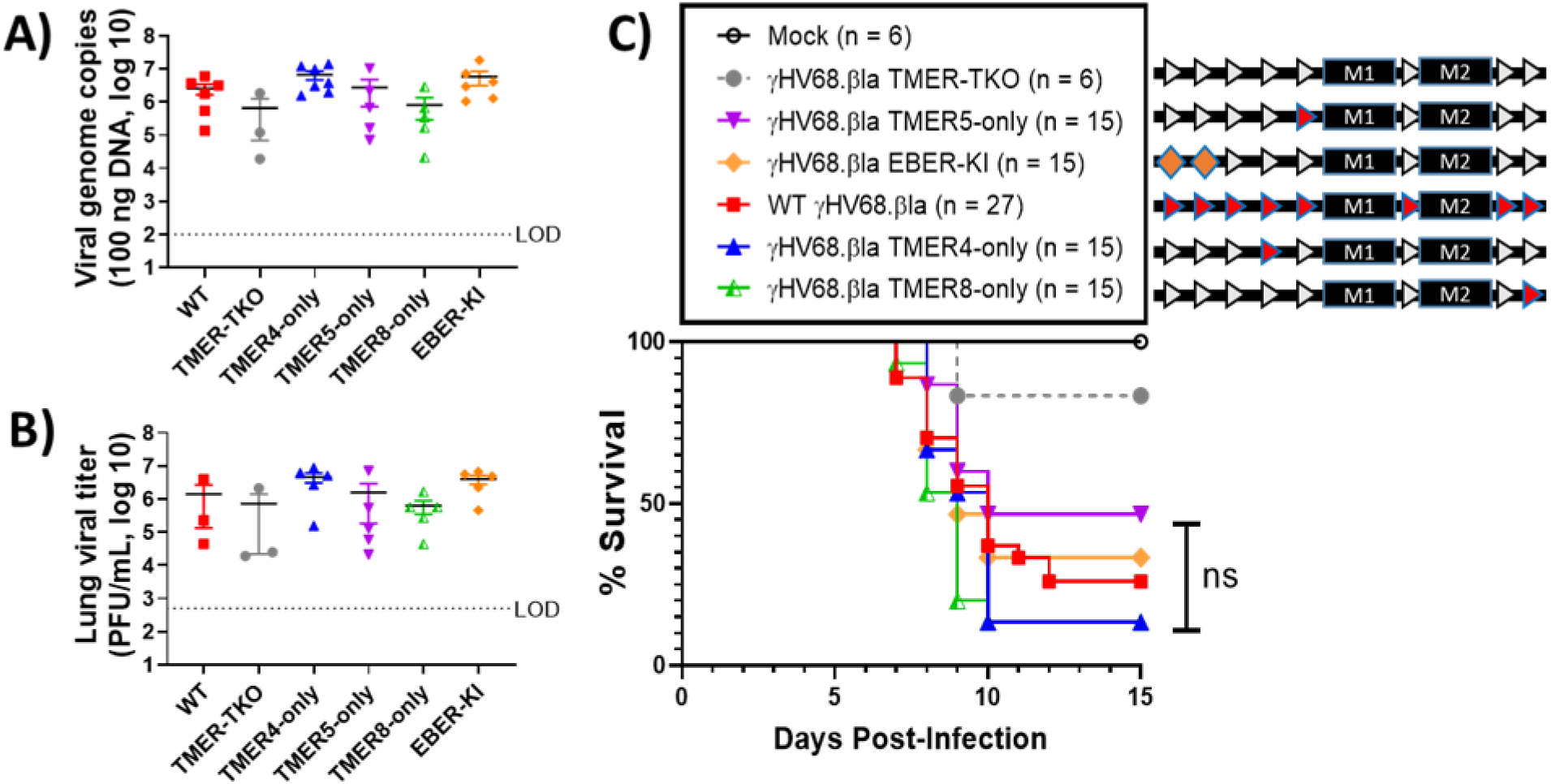
Infection of immunocompromised mice with γHV68 ncRNA recombinants reveal conserved virulence of TMERs and the EBERs. BALB/c IFNγ -/- mice were infected with a panel of γHV68 ncRNA recombinants. At 8 days p.i., lung tissue was collected for viral titer analysis by **(A)** qPCR for viral DNA (gB gene) and **(B)** plaque assay quantitation of infectious virus. Limit of detection (LOD) is indicated by a horizontal dashed line on each graph. Virus was not detected in mock-infected tissue samples in each analysis. Individual symbols represent the value from an individual mouse. Three mice were analyzed for WT and TMER-TKO γHV68, and five mice were analyzed for all other viruses. Horizontal black lines indicate the mean of each group. One-way ANOVA analysis with multiple comparisons of each γHV68 recombinant to WT γHV68 detected no significant difference. **C)** Analysis of BALB/c IFNγ -/- mice following infection with WT or recombinant γHV68 monitored for signs of morbidity over the course of 15 days. The number of mice in each group is indicated. Statistical analysis of survival curves was done by log-rank (Mantel-Cox) test with pairwise comparisons of recombinant viruses and WT γHV68.βla. P-values for survival following infection with each recombinant except TMER-TKO compared to WT virus are all greater than 0.05 (not significantly different, “ns”); TMER-TKO = 0.025, TMER4-only = 0.47, TMER5-only = 0.21, TMER8-only = 0.14, EBER-KI = 0.79.

## DISCUSSION

The γHVs express a diverse set of RNAs that both promote viral propagation and lifelong infection, and actively engage with and manipulate host cell machinery. Among these, the γHV non-coding RNAs, including the EBV EBERS and the γHV68 TMERs, are short non-coding RNAs transcribed by RNA polymerase III. Notably, their expression by RNA pol III likely endows these RNAs with distinct features, including a 5’-triphosphate and 3’-poly U tract, that afford the opportunity of these RNAs to interact with host RNA binding proteins. Indeed, previous studies on the EBV EBERs have demonstrated that these RNAs can engage with both RIG-I, an innate immune sensor that can bind to RNAs containing a 5’-triphsophate, and La, a host RBP that can bind to 3’-poly U tracts (12, 21). While the EBERs and the TMERs are both abundant small, pol III-transcribed ncRNAs, and are predicted to include double-strand RNA segments, these RNAs do not have significant sequence similarity. The TMERs further demonstrate unique characteristics not reported for the EBV EBERs, encompassing bifunctional elements (i.e. a 5’ tRNA-like structure and 3’ microRNAs) and substantial post-transcriptional processing into multiple distinct RNA species (as described in Fig 2 and (8)). Though the viral tRNA-like structures are processed into mature tRNAs with a 3’-CCA addition, they are not aminoacylated, indicating they are very unlikely to directly influence translation (24). Given that the TMERs and EBERs are expressed by two related γHVs that are studied in distinct experimental contexts, whether there are conserved biochemical or functional properties between the EBERs and the TMERs remains poorly understood at this time.

Here, we sought to biochemically characterize the TMERs and determine whether the TMERs and the EBERs share conserved binding properties to the host RBPs, RIG-I and La, and conserved functional properties *in vivo*. To do this, we have leveraged the unique strengths of the γHV68 system, a small animal model of γHV infection and pathogenesis that allows us to study these questions in the context of primary, *de novo* γHV68 infection. Our studies revealed three major findings about the TMERs and EBERs. First, we present evidence that multiple TMER- derived RNAs contain at least a portion of RNAs with a 5’-triphosophate, demonstrated by the sensitivity of certain RNAs to Terminase-mediated degradation only after treatment with RNA- polyphosphatase, building on a previously established method (29). Second, we demonstrate that both the TMERs and EBERs are capable of binding to RIG-I and La during primary infection, but that the strength and/or magnitude of interaction is much greater with the La protein. Third, we report that expression of an individual TMER (TMER 4, 5 or 8), or the EBV EBERs, is capable of restoring the defect in virulence observed in a γHV68 recombinant lacking all 8 TMERs. These studies extend our previous findings that expression of either TMER1, or the tRNA-like portion and partial stem of TMER1, is sufficient to confer virulence. In combination, these studies suggest that miRNA-independent, sequence-diverse features of the TMERs can facilitate pathogenesis in an immune-compromised mouse model (7). The ability of the EBV EBERs to restore virulence is consistent with a recent report of an independent single EBER- knock-in γHV68 recombinant capable of restoring certain deficits of primary infection observed when using a virus that lacks certain TMERs (35). More broadly, these studies suggest that multiple γHV ncRNAs possess a functionally conserved ability to enhance *in vivo* pathogenesis in an immune-compromised mouse model (7), a property that may be linked to conserved interactions with host RBPs.

Although our data demonstrate that the EBERs and TMERs have certain conserved biochemical and functional properties, it is important to acknowledge that this does not mean that individual viral ncRNAs don’t have unique functions as well. For the EBV EBERs, multiple reports have demonstrated that EBER1 and EBER2 have unique binding partners and functions are not simply redundant elements (36–38). For the TMERs, it is notable that each of the 8 TMERs encodes 1-2 distinct miRNAs, with a range of targets (27, 39). In addition, mutagenesis studies have identified a specific role for TMER4 in promoting hematogenous dissemination of γHV68 during primary infection *in vivo* (35), a function that can be restored by EBER1 but not another viral pol III-transcribed ncRNA, adenovirus VAI (40). These studies, in combination with data presented here demonstrating conserved biochemical and functional characteristics, strongly suggest that these viral ncRNAs make both unique and redundant contributions to enhance γHV infection. We anticipate that the continued analysis of the viral ncRNA recombinants presented here will enable new insights into conserved and functionally redundant properties of these ncRNAs, and will complement ongoing studies in which individual ncRNAs are specifically disrupted in the context of the virus (40–42).

Our studies raise a number of important future questions. *First, what are the conserved functional properties and targets of the TMERs or EBERs that promote in vivo pathogenesis?* Our studies implicate possible pol III-RNA associated features (e.g. 3’-poly U tract) as one potentially conserved mechanism by which these ncRNAs may engage host pathways and influence the host inflammatory or immune response. A top candidate in this regard is the conserved binding of the TMERs and EBERs to La, a condition we observed under stringent wash conditions. Studies on the EBERs have indicated that EBER/La complexes can serve as TLR3 ligands, to engage the innate immune response (10, 11, 22). Whether this process occurs during γHV68 infection, and what genetic contribution this putative pathway has on the outcome of infection remains an important and unresolved question that is uniquely capable of being addressed using the γHV68 system, leveraging our panel of ncRNA recombinants. Another important property of the TMERs and the EBERs that cannot be ignored is that these RNAs are extremely abundant, potentially altering the host cell by overwhelming or outcompeting host processing machinery and innate immune sensing pathways (32, 43). In this later model, modulating or antagonizing viral ncRNA abundance would be predicted to blunt the effects of these ncRNAs, a potential future therapeutic opportunity. *A second major question that these studies raise is what impact, if any, does cellular context and state of infection (i.e. lytic vs. latent infection) have on ncRNA/host RBP interactions and ncRNA function?* Here, our studies focused on primary *de novo in vitro* infection of fibroblasts and *in vivo* infection, in which multiple cell types are lytically infected with little to no latent infection. Whether viral ncRNA/host RBP interactions vary as a function of cell type or during lytic versus latent infection is a vital question to understand the complexity of host engagement by these ncRNAs, and will likely reveal context-specific regulation of these ncRNAs and their interactions.

In conclusion, our studies on two distinct classes of γHV ncRNAs, the TMERs and EBERs, demonstrate that these ncRNAs have conserved biochemical and functional features, including the ability to interact with the host RNA-binding protein, La, and to enhance virulence in an *in vivo* model of γHV pathogenesis. The conserved properties of these RNAs despite significant sequence divergence strongly supports the concept that a major biological function of these ncRNAs is mediated either through conserved biochemical or structural features or as a consequence of the unique abundance of these viral RNAs, expressed throughout the viral life cycle. We anticipate that future studies leveraging the unique strengths of this experimental system will allow us to critically assess both unique and conserved features of individual viral ncRNAs *in vitro* and *in vivo*.

## METHODS

### Viruses and tissue culture

Human endothelial kidney (HEK 293) and murine fibroblast 3T12 cells (ATCC CCL-164) were cultured in Dulbecco’s modified Eagle medium (DMEM; Life Technologies) supplemented with 5% fetal bovine serum (FBS; Atlanta Biologicals), 2 mM L-glutamine, 10 U/ml penicillin, and 10 μg/ml streptomycin sulfate (complete DMEM, cDMEM). Vero-cre cells (gift from David Leib at Dartmouth School of Medicine) were grown in complete DMEM with 10% FBS. Cells were cultured at 37°C with 5% CO2.

All viruses and recombinants were derived from γHV68 strain WUMS (ATCC VR-1465) (44). Mutants were created using a BAC harboring wild-type γHV68 (45) or γHV68.ORF73βla (referred to subsequently as γHV68.βla) (46). TMER-TKO and TMER1-only γHV68.βla were previously characterized (7), and new γHV68 recombinant viruses were generated via *en passant mutagenesis* (47). The γHV68.βla TMER4-only, TMER5-only, TMER8-only, and γHV68.βla EBER-knock in mutants were generated by Gibson assembly of two synthetic DNA fragments (gBLOCKS: New England Biolabs) followed by PCR and subsequent electroporation of the assembled and PCR-amplified fragments into E. coli strain GS 1783.5 harboring the γHV68.βla BAC. The presence of the individual TMERs and the EBER knock-in was verified by sequencing across the locus. For the EBER-knock in mutant, we replaced 563bp of the WT γHV68 TMER 1-2 region with 502 bp of the EBER 1-2 sequence (nt 127-629 of WT TMER sequence in γHV68.βla; nt 6629-7130 of EBV Genbank AJ507799.2). Recombinants were confirmed by restriction digestion of BAC DNA and PCR products from the left end of the γHV68 genome, as well as sequencing of the left end of the γHV68 genome using the primers listed in Table 2.

**Table 2.**
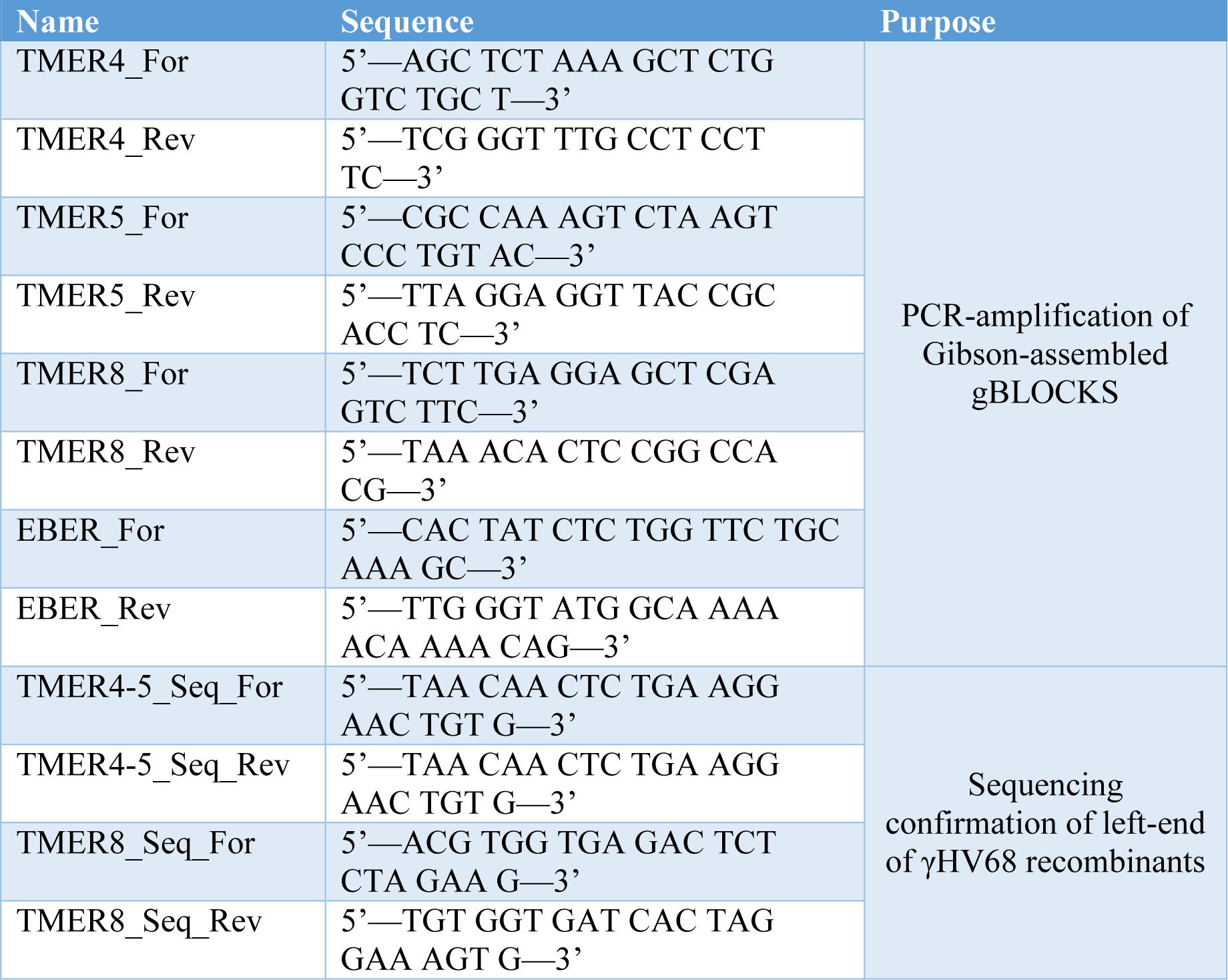
Primers used for recombinant gBLOCK assembly and sequencing of γHV68 recombinants.

Infectious virus was generated from confirmed BACs by transfecting BAC DNA into HEK 293 cells. Resulting virus was passaged through cre-recombinase expressing Vero cells to remove the loxP-flanked BAC origin of replication (48). Removal of the BAC origin of replication from the viral DNA was confirmed by PCR analysis (not shown).

### Infections

To ensure an accurate count, and multiplicity of infection, all infections were done using fresh cell counts, determined by removing one well of cells with 0.05% Trypsin-EDTA (Life Tech, Cat. No. 25300-054), mixing with Trypan Blue dye (Bio-Rad, Cat. No. 145- 0021), and counting live cells with the TC20 Automated Cell Counter (Bio-Rad). Virus stock titers were previously quantified with at least three plaque assays. The appropriate amount of viral stock was mixed with 5% FBS DMEM to make 100 μL (12-well format) or 250 to 300 μL (6-well format) of viral inoculum per well at a multiplicity of infection (MOI) of 5. Cells were incubated at 37°C with 5% CO_2_ for 1 hour and rocked every 15 min before viral inoculum was removed. Cells were rinsed with PBS and covered with 2 mL of complete 5% DMEM. Samples were harvested at the indicated hours post-infection (hpi).

### 5’-end characterization of small RNAs

Characterization of the 5’ ends of RNAs was performed as previously described (29); however, we analyzed all RNA species under 300 nucleotides rather than 70 nucleotides (29) - a generous cutoff to ensure inclusion of all TMER gene products. Briefly, HEK 293 cells were mock treated or infected with either WT or TMER1- only γHV68.βla at a MOI of 5. RNA was isolated from cells 24 hpi using PIG-B (49) and chloroform/isopropanol extraction, then fractionated on 15% polyacrylamide gel electrophoresis (PAGE)-urea gel. All RNA species smaller than 300 nucleotides were excised in roughly 5 mm² sections and dissolved in 1M NaCl at 4°C for approximately 48 hours. Gel pieces were removed from RNA samples with a 100 μM strainer (Fisher, Cat. No. 22363549), then RNA was concentrated with Sartorius Vivaspin 15R columns (Sartorius, Cat. No. VS15RH92) and precipitated with isopropanol and 0.5M final concentration of ammonium acetate. Isolated small RNAs (4 to 5 μg) were incubated with 1 μL (for 4 μg RNA), 1.5 μL (for 4.5 μg), or 2 μL (for 5 μg RNA) of RNA 5’-Polyphosphatase (Epicentre, Cat. No. RP8092H; 20 units/μL) or no enzyme in a 20 μL reaction for 1 hour at 37°C. RNA was then precipitated and remaining small RNAs (up to 1 μg) were incubated with 1 μL Terminator™ 5’-Phosphate-Dependent Exonuclease (Epicentre, Cat. No. TER51020; 1 unit/μL) or no enzyme in a 20 μL reaction for 3 hours at 30°C. Each RNA 5’-Polyphosphatase and Terminator™ reaction included 1.5 or 0.5 μL RNase inhibitor (NEB, Cat. No. M0307L; 40 units/μL), respectively. Resulting RNA samples were analyzed by northern blot.

### Northern Blot

Northern blots were performed as previously described (8, 50). Enzyme-treated RNA samples were run on a 12% denaturing PAGE-urea gel, transferred to Zeta-Probe® GT Blotting Membrane (Biorad, Cat. No. 1620196), and detected with the indicated RNA oligonucleotide probes and conditions (Table 1). Relative density of each band on the blotting membrane was determined by normalizing to ethidium bromide-stained 5S rRNA in the PAGE-urea gel using ImageJ software (51). Band density was calculated as a fold change of the related untreated RNA population, which was set to 1.

### Transfecting and Infecting Cells

For transfections, HEK 293 cells were cultured in 5% FBS DMEM without penicillin or streptomycin for approximately 24 hours. Cells were plated in 2 mL at 2×10^5^ cells per mL in 6-well plates. Transfection solution for each well contained Opti-MEM (Thermo Fisher Scientific) and 2 μg of pEFBos FLAG-RIG-I (Gale Lab), pCMV2-human FLAG-La (Maraia lab), or pcDNA5 FRT/TO FLAG-HA GFP (Gack lab). Solutions for experiments using plasmid-expressed EBERs included 2 μg of pSP73-EBERs plasmid (Steiz lab) for a total of 4 μg of plasmid DNA. After plasmids were added to the Opti-MEM for a total volume of 200 μL, solutions were incubated with 4:1 X-tremeGENE HP DNA Transfection Reagent (Sigma-Aldrich) for at least 15 min at room temperature. Transfection solution was added drop-wise to cells, then cells were incubated at 37°C in 5% CO_2_ overnight. HEK 293 cells were infected 24 hours post-transfection at an MOI of 5 as previously described.

### Immunoprecipitation

Cell lysates were harvested 24 hpi by scraping and pelleting cells and supernatants at 500xg for 10 min. 6 wells were pooled into one 15 mL conical tube for each sample. Cell pellets were rinsed in cold PBS following removal of supernatant, transferred to 2 mL screwcap tubes, then pelleted again as before. PBS was removed from the cell pellet and cells were resuspended in 500 μL of lysis buffer. Lysis buffer contained 50 mM Tris pH 7.4, 150 m NaCl, 1% NP-40, dithiothreitol (DTT), sodium fluoride (NaF), and protease inhibitors (aprotinin, leupeptin, and phenylmethylsulfonyl fluoride [PMSF]). Following incubation with lysis buffer, cell lysates were cleared by centrifugation at 10,000xg for 20 min at 4°C and transferred to new 2 mL screw cap tubes. Protein concentrations of cell lysates were determined using the Pierce BCA Protein Assay kit (Thermo Fisher Scientific, Cat. No. 23227).

350 μg of cell lysates were incubated at 4°C overnight on a rotator with anti-FLAG M2 magnetic beads (Sigma, Cat. No. M8823) and TBS (50 mM Tris pH 7.4, 150 mM NaCl). In permissive wash conditions, beads were washed three times with cold TBS (Fig 3, Fig S3). In stringent wash conditions, beads were washed three times with cold lysis buffer (with DTT, NaF, and protease inhibitors), then two times with cold high-salt lysis buffer (50mM Tris-HCl pH 7.4, 300mM NaCl, 1% NP-40, DTT, NaF, protease inhibitors). Following removal of wash buffer, beads were resuspended in 60 μL of storage buffer (50mM Tris pH 7.4, 150mM NaCl, 50% glycerol, 0.02% Na Azide) and 10% of the volume was transferred to a new tube for western blot analysis of proteins. The remaining volume was used for RNA isolation.

### Western Blot

Total cell lysate (35 μg) or immunoprecipitation beads were incubated with 4x Laemmli buffer (52) at 95°C for 5 min, then loaded into a 10% or 12% SDS-PAGE gel with Precision Plus Protein Dual Color Standard (Biorad, Cat. No. 1610374). Following adequate separation of bands, proteins were semidry transferred to Immobilon®-P PVDF membrane (MilliporeSigma, Cat. No. IPVH00010) for 1 h to 1 h 15 min at 10 Volts (Thermo Fisher Scientific, Owl™ HEP-1). Membrane was probed with 1:2000 M2 monoclonal mouse anti-FLAG (Sigma, Cat. No. F1804-50ug). Proteins were detected with horseradish peroxidase (HRP)-conjugated donkey anti-mouse secondary antibody (Jackson ImmunoResearch, Cat. No. 715-035-150) and Amersham™ ECL™ Prime Western Blotting Detection Reagent (Cytiva, Cat. No. RPN2232).

### RNA analysis

#### Isolation

RNA was isolated from immunoprecipitation complexes or cells with TRIzol® Reagent (Thermo Fisher Scientific, Cat. No. 15596026), then DNase-treated with TURBO^TM^ DNase (Invitrogen, Cat. No. AM2238) following the manufacturer’s protocols.

#### Reverse transcription PCR amplification

Primers used for RT-PCR are listed in Table 3. RNA transcripts of interest were detected using the OneStep RT-PCR Kit (Qiagen, Cat. No. 210212), with the following conditions: (i) 50°C for 30 m, (ii) 95°C for 15 m, (iii) 40 cycles of 94°C for 30 s, anneal for 30 s, and 72°C for 30 s, (iv) 72°C for 10 m, (v) hold at 4°C. The annealing temperature was adjusted depending on the target: M3 = 50°C, TMER1 = 50.5°C or 52°C (for Fig 3 and Fig 5, respectively), TMER4 = 50°C, TMER5 = 50°C, TMER8 = 56°C, EBER1 = 52°C, EBER2 = 56°C. For semi-quantitative analysis, step (iii) was limited to 30 cycles (TMER1, EBER1, EBER2) or 33 cycles (TMER4).

**Table 3:**
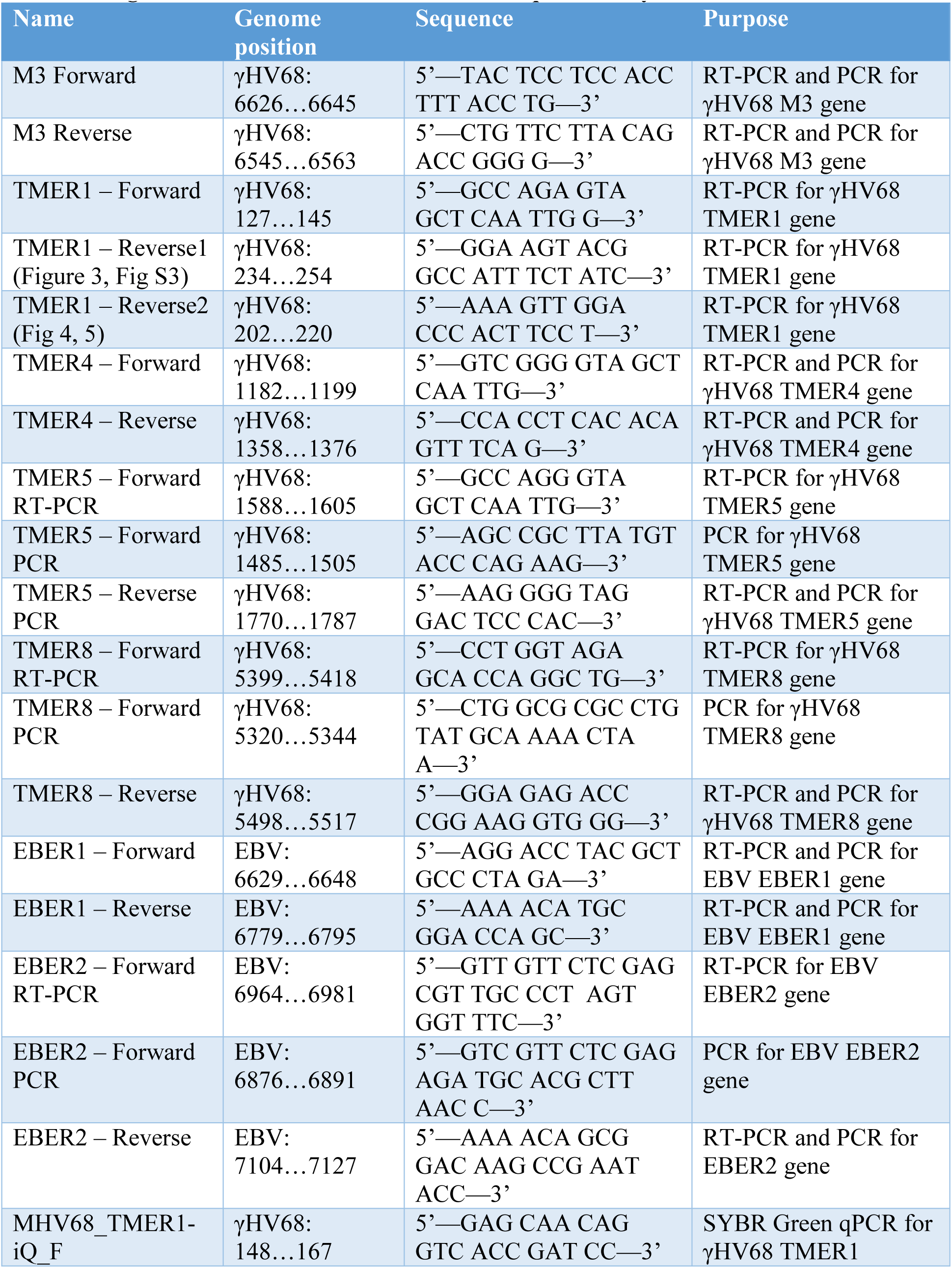
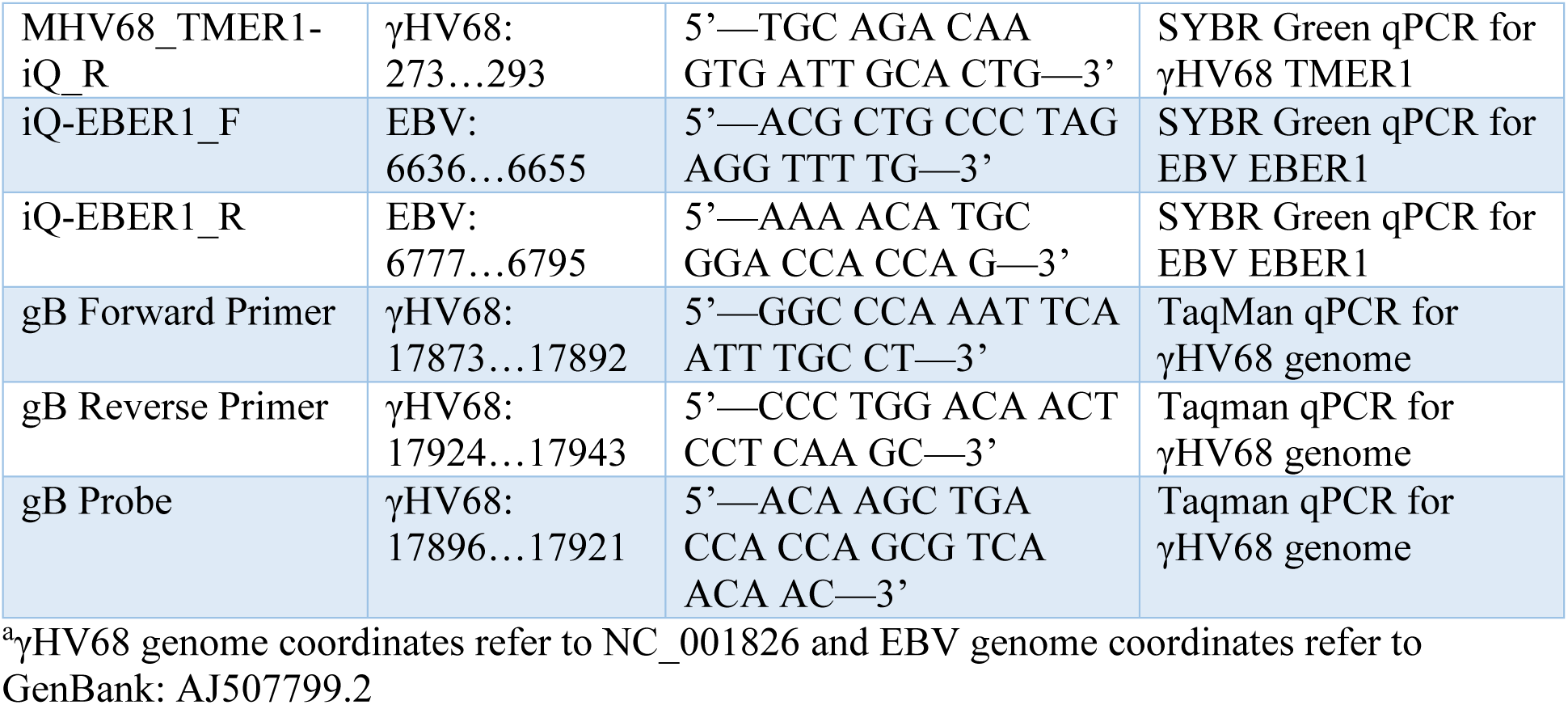
Oligonucleotides for RT-PCR, PCR, and qPCR analysis.

#### PCR analysis

PCR amplification was performed using Taq DNA polymerase (Qiagen, Cat. No. 201205) with the following conditions: (i) 95°C for 5 min, (ii) 30-40 cycles of 94°C for 30 s, anneal for 30 s, 72°C for 30 s, (iii) 72°C for 10 min, (iv) hold at 4°C. When checking RNA for DNA contamination, the same primers as the RT-PCR reaction were used, and step (iii) annealing temperature and cycle numbers were adjusted for consistency with the related RT-PCR reaction. For DNA targets, primers are listed in Table 3 and the annealing temperature was adjusted depending on the target: M3 = 50°C, TMER4 = 50°C, TMER5 = 53°C, TMER8 = 60°C, EBER1 = 52°C, EBER2 = 56°C.

#### Reverse transcription quantitative PCR

RNA samples shown to be DNA-free by PCR were converted to cDNA using SuperScript III Reverse Transcriptase (Invitrogen, Cat. No. 18080093) following the manufacturer’s protocol. 20 nanograms (ng) of the cDNA was then used for qPCR analysis of the γHV68 TMER1 (31) or EBV EBER1 genes using iQ™ SYBR® Green Supermix (Bio-Rad, Cat. No. 1708880) with the following conditions: i) 95°C for 3 min, ii) 40 cycles of 95°C for 15 s, 60°C for 1 min, iii) 95°C for 15 s, 60°C for 1 min, 95°C for 15 s. Primers used for qPCR analysis are listed in Table 3.

#### RT-qPCR analysis

Viral ncRNAs were detected by qPCR from whole cell lysates (“total”) and immunoprecipitated (“IP”) proteins (GFP, RIG-I, or La) in both mock and infected (WT or EBER-KI γHV68.βla) conditions. The Ct value for each IP sample was subtracted from the Ct value for the matched total sample; (𝐶𝑡_𝑝𝑟𝑜𝑡𝑒𝑖𝑛 𝑡𝑜𝑡𝑎𝑙_ − 𝐶𝑡_𝑝𝑟𝑜𝑡𝑒𝑖𝑛 𝐼𝑃_). This value was then normalized to the GFP control by subtracting from the matched GFP condition for the “delta Ct” (ΔCt). For RIG-I samples: 𝛥𝐶𝑡 = (𝐶𝑡_𝑅𝐼𝐺−𝐼 𝑡𝑜𝑡𝑎𝑙_ − 𝐶𝑡_𝑅𝐼𝐺−𝐼 𝐼𝑃_) − (𝐶𝑡_𝐺𝐹𝑃 𝑡𝑜𝑡𝑎𝑙_ − 𝐶𝑡_𝐺𝐹𝑃 𝐼𝑃_). For La samples: 𝛥𝐶𝑡 = (𝐶𝑡_𝐿𝑎 𝑡𝑜𝑡𝑎𝑙_ − 𝐶𝑡_𝐿𝑎 𝐼𝑃_) − (𝐶𝑡_𝐺𝐹𝑃 𝑡𝑜𝑡𝑎𝑙_ − 𝐶𝑡_𝐺𝐹𝑃 𝐼𝑃_). Fold enrichment for each target was calculated as 2^Δ𝐶𝑡^.

#### Single-step replication analysis

3T12 fibroblasts cells were infected in triplicate with each viral recombinant at an MOI of 5 for single-step replication analysis as previously described (53). Cells were inoculated with virus for 1 h at 37°C in 5% CO_2_, rinsed with PBS, then incubated in 5% cDMEM. One set of samples (cells and supernatant) was collected immediately following inoculation (0 hpi). Remaining samples were collected at 1, 6, 12, 24, and 48 hpi, then subjected to three freeze-thaw cycles before plaque assay quantitation (33).

### *In vivo* infections

#### Mice

BALB/c interferon-gamma deficient mice (IFNγ^-/-^) were originally obtained from the Jackson Laboratory [strain C.129S7(B6)-Ifngtm1Ts/J, stock no. 002286]. Mice were bred in- house at the University of Colorado Denver Anschutz Medical Campus following university regulations and infected mice were housed in an animal biosafety level 2 facility in accordance with all university regulations.

#### Virus infection of mice

Mice were anesthetized with isoflurane (McKesson, Cat. No. 803250) and intranasally inoculated with 4×10^5^ PFU/mouse of WT γHV68.βla or the indicated viral recombinants in a 40-μl total volume of 5% cDMEM as previously described (7). Mice were monitored daily for signs of disease. Any mice that appeared moribund were sacrificed, and their lungs were removed for viral titer analysis.

#### *Ex vivo* viral titer analysis

Infected tissues were collected at the indicated time post-infection and frozen at -80°C. The right post-caval lobe of the lung was separated for qPCR and plaque assay analysis. DNA was isolated from a portion of the lung tissue using the Qiagen DNeasy® Blood & Tissue Kit (Cat. No. 69506) with a modified overnight proteinase K incubation followed by heat inactivation (95°C for 10 min). Isolated DNA was subsequently analyzed by qPCR for a viral gene (gB). Virus titer was quantified in the remaining lung tissue by homogenizing tissue in 1 mL of 5% cDMEM and 1.0-mm silica beads (BioSpec Products Inc, Cat. No. 11079110z) via MagNA Lyser (Roche). Homogenized tissue was subjected to three freeze-thaw cycles prior to plaque assay quantification of viral titer (53).

#### Quantitative reverse-transcription PCR for viral genome

The number of viral genome copies in DNA samples was quantified by qPCR for the viral gene gB. Lung DNA was normalized to a concentration of 20 ng/μL for qPCR analysis of 100 ng of DNA using a LightCycler 480 probe kit (Roche, Cat. No. 04707494001) with the gB primers and probe listed in Table 3 (54, 55). A gB standard curve was generated using a gB plasmid dilution series ranging from 10^2^ copies to 10^10^ diluted in background DNA, with a limit of detection (LOD) of 100 copies (56). Each sample was run as a technical triplicate.

#### Plaque Assay

Plaque assay quantification of viral titer was performed as previously described (53, 57) with the following modifications. 3T12 fibroblasts were plated in 12-well plates at 8.5×10^4^ cells per well the day prior to infection. Viral samples were diluted 10-fold in 5% cDMEM, then 100 μL of inoculum was added to each well. An internal standard was included for each infection to ensure reproducible sensitivity for each plaque assay. Cells were incubated with virus for 1 h at 37°C at 5% CO_2_ and plates were rocked every 15 min. Cells were then overlaid in a 1:1 mix of 10% cDMEM and carboxymethyl cellulose (CMC; Sigma, Cat. No. C- 4888) supplemented with Gibco™ Amphotericin B (Thermo Fisher Scientific, Cat. No. 15290018). 8 days post infection, the overlay was removed and cells were rinsed with PBS. Cells were then stained with 0.5% methylene blue in 70% methanol and plaques were counted to calculate viral titer.

#### Software and Statistical Analysis

Analysis of northern blot images and band densities were performed using ImageJ software (51). Plotting and data analysis were performed using GraphPad Prism (version 9.3.1; GraphPad Software, San Diego, California USA, www.graphpad.com). Statistical significance was tested by one-way or two-way analysis of variance (ANOVA) for comparing three or more conditions or comparing grouped data, respectively.

#### Ethics Statement

All animal studies were performed in accordance with the recommendations in the Guide for the Care and Use of Laboratory Animals of the National Institutes of Health.

Studies were conducted in accordance with the University of Colorado Denver Institutional Animal Use and Care Committee under the Animal Welfare Assurance of Compliance policy (no. D16-00171). All procedures were performed under isoflurane anesthesia, and all efforts were made to minimize suffering.

## ACKNOWLEDGEMENTS

This research was funded by NIH R21AI134084, R01AI121300, R01AI157201 and the Molecular Pathogenesis of Infectious Disease Training Grant 5T32AI052066.

We thank the members of the van Dyk and Clambey lab for helpful discussions and members of the Colorado RNA Bioscience Initiative for their insights. Special thanks go to Dr. Neelanjan Mukherjee for expert advice on IP experiments. We thank the following labs for their generous plasmid gifts: Dr. Michael Gale (pEFBos FLAG-RIG-I), Dr. Rich Maraia (pCMV2- human FLAG-La), Dr. Joan Steitz (pSP73-EBERs), and Dr. Michaela Gack (pcDNA5 FRT/TO FLAG-HA GFP).

---

**Figure S1.**
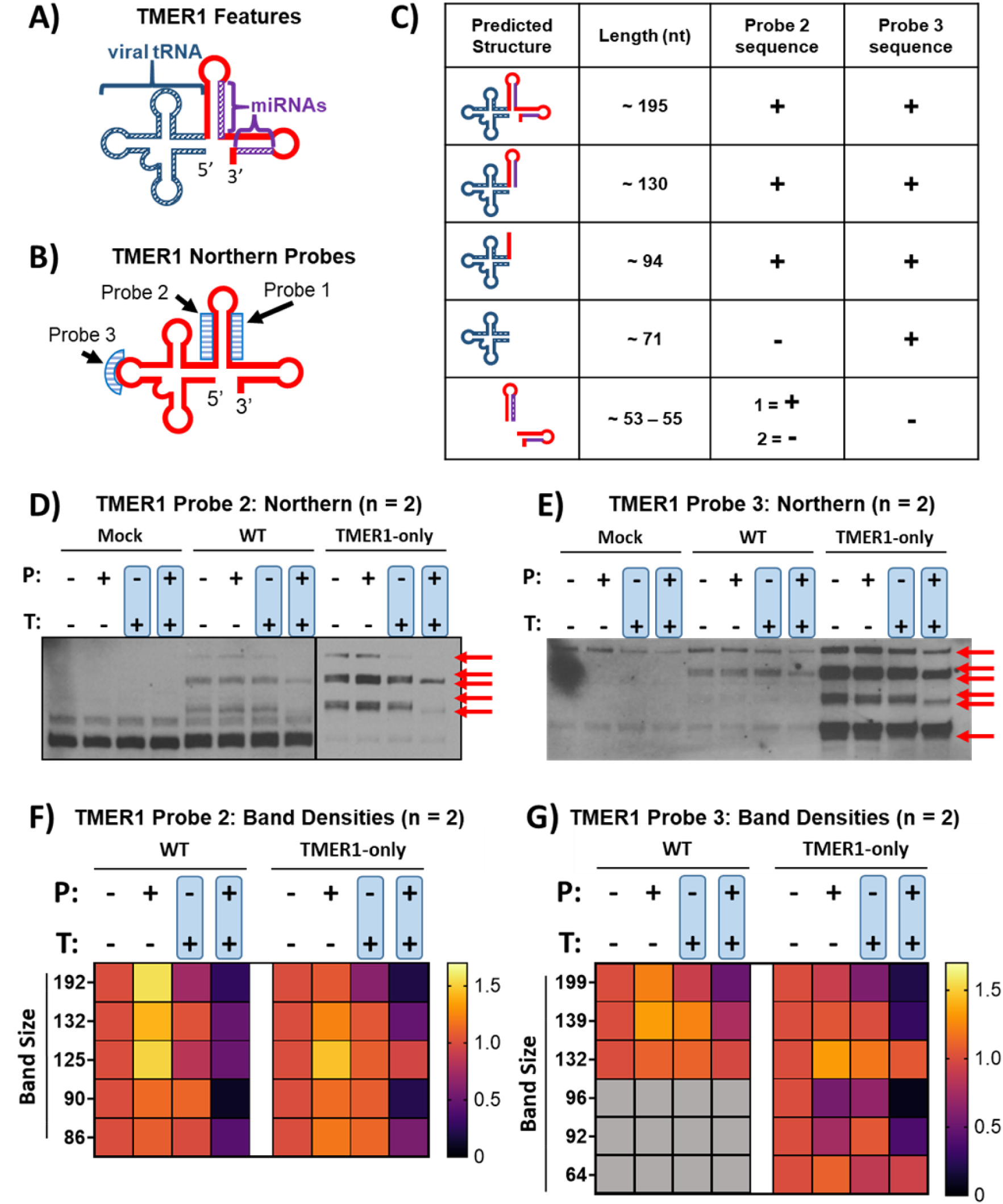
Multiple distinct TMER probes detect 5’-triphosphate on TMER1 RNAs. **A)** Schematic of features of TMER1 RNA. The TMER1 predicted structure consists of a tRNA-like loop (dark blue) and multiple stem loops that are processed into biologically active miRNAs (purple). **B)** Schematic of northern probe sequences used to detect TMER1. Different northern probes (light blue boxes) bind to various regions of TMER1 RNA, allowing detection of alternate, processed forms. **C)** Table showing the multiple possible alternate forms of TMER1 with varying lengths. The two probe sequences shown here (Probe 2 and Probe 3) are present in some TMER1 forms (+), but not others (-). Following sequential enzymatic treatments (P = RNA 5’-polyphosphatase, T = Terminator™), small RNAs were resolved by SDS-PAGE gel and northern blot was performed with TMER1 Probe 2 **(D)** or Probe 3 **(E).** The RNA bands specific to TMER1 are marked with red arrows. Band densities for the TMER1 RNAs detected by TMER1 probe 2 **(F)** and TMER1 probe 3 **(G**). Densities of the TMER1 northern blot bands were normalized to a 5S rRNA loading control stained with ethidium bromide. Relative density of bands were calculated as a fold change of the untreated RNA population, which was set to 1, and presented as heat maps. Band sizes were calculated as averages based on migration of a ladder included in each experiment. RNA bands not consistently detectable in WT γHV68 infection are shown as gray boxes. Data for each probe is from two independent experiments.

**Figure S2.**
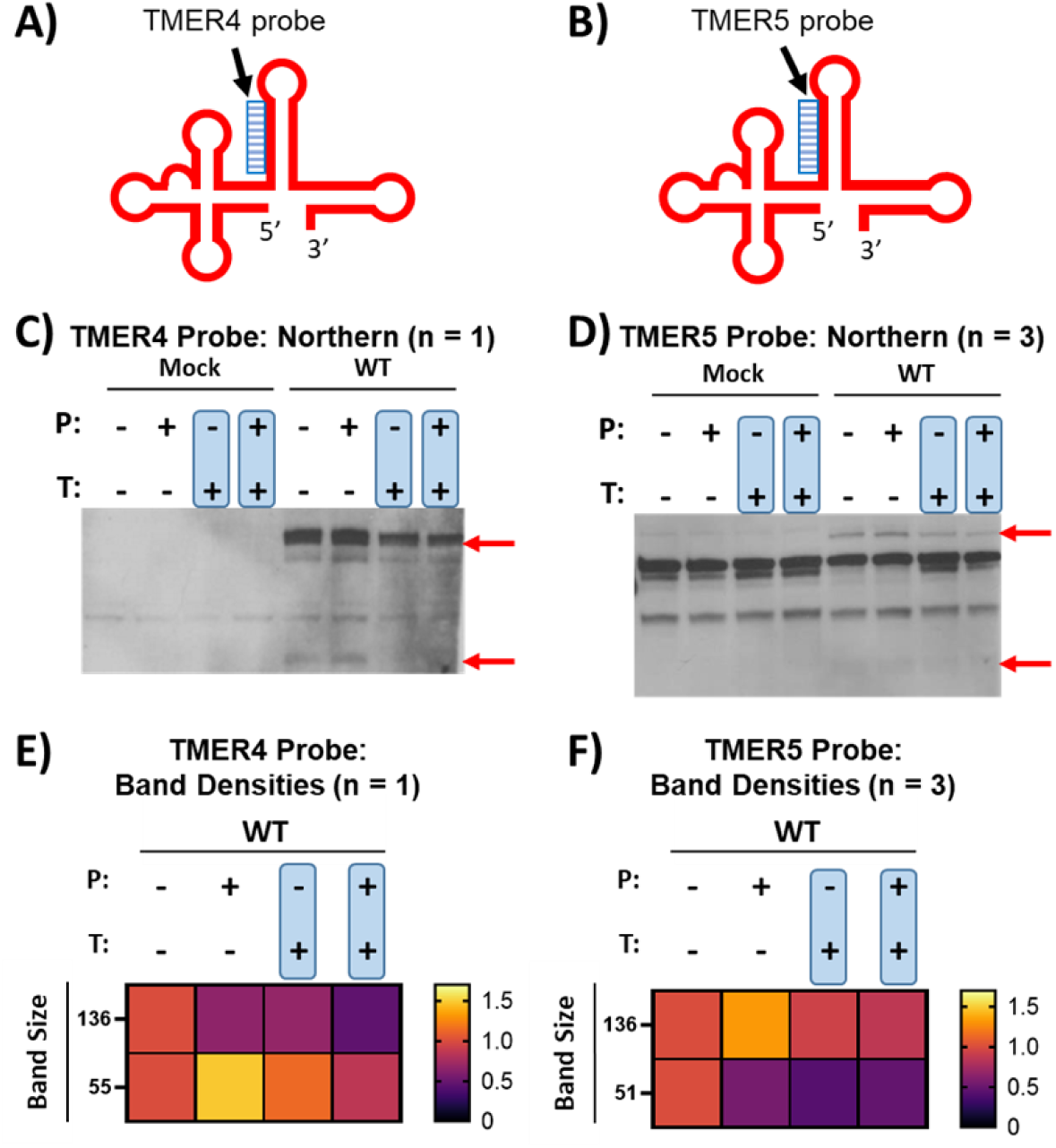
5’ RNA end characterization of TMER4 and TMER5. Predicted secondary structures for TMER4 **(A)** and TMER5 **(B).** The sequence for northern probes used to detect each TMER (light blue boxes) is present in the 5’ end of the first stem loop of the individual TMERs. Following sequential enzymatic treatments (P = RNA 5’- polyphosphatase, T = Terminator™), small RNAs were resolved by SDS-PAGE gel and northern blot was performed with probes for TMER4 **(C)** or TMER5 **(D).** The RNA bands specific to TMER4 or TMER5 are marked with red arrows. Band densities for the RNAs detected by probes targeting TMER4 **(E)** and TMER5 **(G**). Densities were normalized as previously described and presented as heat maps. Analysis was performed in one experiment (TMER4) or three independent experiments (TMER5).

**Figure S3.**
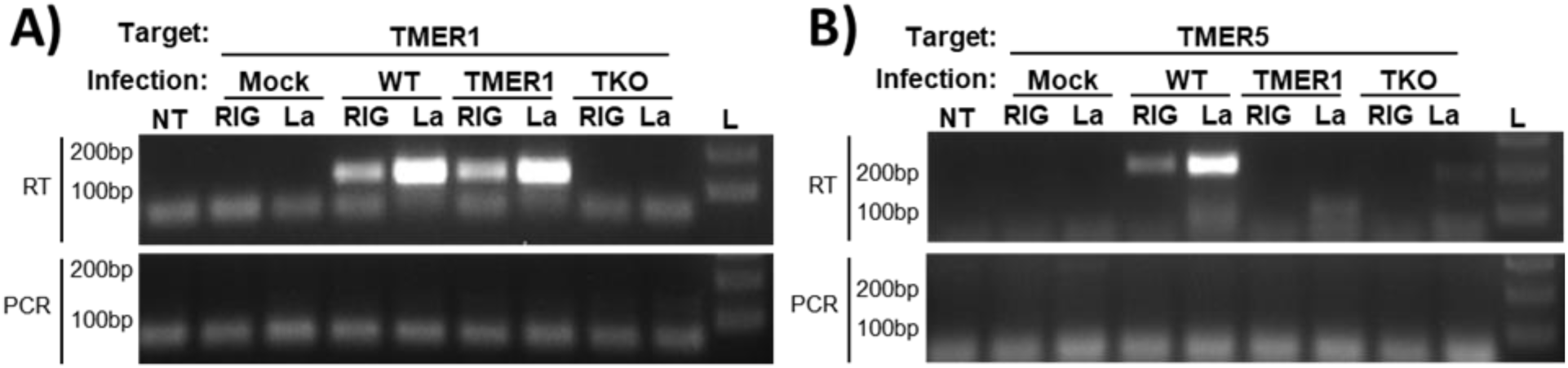
Immunoprecipitation of FLAG-tagged proteins indicates multiple TMERs interact with host proteins. HEK 293 cells were transfected with FLAG-tagged RIG-I or La, then infected with WT, TMER1-only (TMER1), or TMER-TKO (TKO) γHV68 as previously described. Whole cell lysates were collected 24 hpi and used for immunoprecipitation of FLAG-tagged RIG-I or La. RNA was isolated from immunoprecipitated complexes and analyzed by RT-PCR with primers targeting TMER1 **(A)** or TMER5 **(B)**. PCR without reverse transcription (“PCR”) was performed in conjunction with RT-PCR to test for DNA contamination. NT = non-template control, L = ladder. Data are representative of one experiment with technical triplicates (TMER1) or duplicates (TMER5).

